# A single-cell transcriptomic atlas characterizes liver non-parenchymal cells in healthy and diseased mice

**DOI:** 10.1101/2021.07.06.451396

**Authors:** Zheng Wang, Jingyang Qian, Xiaoyan Lu, Ping Zhang, Rongfang Guo, He Lou, Shuying Zhang, Jihong Yang, Xiaohui Fan

**Affiliations:** Pharmaceutical Informatics Institute, College of Pharmaceutical Sciences, Zhejiang University, Hangzhou, 310058, China; State Key Laboratory of Modern Chinese Medicine, Tianjin University of Traditional Chinese Medicine, Tianjin 301617, China; Department of Medicine, Columbia Center for Human Development, Columbia University Irving Medical Center, New York, NY, 10032, USA; iMedicine Lab, Alibaba-Zhejiang University Joint Research Center of Future Digital Healthcare, Hangzhou 310058, China

## Abstract

The heterogeneity of liver non-parenchymal cells (NPCs) is essential for liver structure and function. However, the current understanding of liver NPCs, especially in different liver diseases, remains incompletely elucidated. Here, a single-cell transcriptome atlas of 171,814 NPCs from healthy and 5 typical liver disease mouse models, including alcoholic liver disease, nonalcoholic steatohepatitis (NASH), drug-induced liver injury, cholestatic, and ischemia-reperfusion liver injury is constructed. The inter- and intra-group heterogeneity of 12 types (and numerous subtypes) of NPCs involving endothelial cells, hepatic stellate cells (HSCs), neutrophils, T cells, and mononuclear phagocytes (MPs) are summarized. A protective subtype of neutrophils characterized by *Chil3^high^* is validated and found significantly increasing only in drug-induced and cholestatic liver injury models. Transcriptional regulatory network analysis reveals disease-specific transcriptional reprogramming. Metabolic activity analysis indicates that fibrosis is accompanied by increases in glycolysis and retinol metabolism in activated HSCs and MPs. Moreover, we found that cell-cell interactions between cholangiocytes and immune cells contribute more to cholestatic liver fibrosis compared with NASH, while HSCs are more important for NASH fibrosis. Our atlas, together with an interactive website provides a systematic view of highly heterogeneous NPCs and a valuable resource to better understand pathological mechanisms underlying liver diseases.

## INTRODUCTION

The liver is a complex ecosystem, composed of diverse types of cells, that plays vital metabolic and immunological functions (1). Despite considerable improvements over past decades, liver diseases remain a major public health challenge worldwide. Alcoholic liver disease (ALD), nonalcoholic fatty liver disease (NAFLD), drug-induced liver injury (DILI), cholestatic liver injury, and liver ischemia-reperfusion (IR) injury caused by surgery together account for over 70% of the incidence of liver diseases, seriously affecting the quality of human life (2). A major obstacle for development of precision therapies for liver disease is our lack of systematical understanding of the ecosystem, especially in different liver diseases.

Almost all types of liver disease are accompanied by an inflammatory response (3). Liver non-parenchymal cells (NPCs), including mononuclear phagocytes (MPs), endothelial cells (ECs), hepatic stellate cells (HSCs), cholangiocytes, and other infiltrated inflammatory cells (e.g. neutrophils), are essential for the liver structure, function, and response to inflammatory liver injury (4). MPs are composed of Kupffer cells (KCs), monocyte-derived macrophages (MoMFs), and dendritic cells (DCs) (4). KCs, resident macrophages, play a key role in liver inflammation. After activation, KCs adopt M1-like pro-inflammatory macrophage or M2-like anti-inflammatory macrophage functions in response to liver injury (3). Recruited MoMFs, which are divided according to pro-inflammatory (M1) and wound-healing (M2) phenotypes, also play a role in acute and chronic liver inflammation (3). Although KCs and MoMFs have similarities, they can be distinguished by numerous markers (3). Neutrophils also play essential roles in acute and chronic inflammation (5). These inflammatory cells also have interaction with HSCs through specific ligand-receptor pairs (6). Inflammatory activities of these inflammatory cells induce HSCs activation (from the resting phenotype to a myofibroblast-like phenotype), which is the major cause of liver fibrosis (7). Activated HSCs themselves promote further liver inflammation and fibrosis, which is characterized by increased cell proliferation, the secretion of pro-inflammatory cytokines, and an enhancement in the synthesis of extracellular matrix (ECM) (3). Thus, liver NPCs show considerable cellular diversity, and their crosstalk plays an important role in liver disease. Although it is well known that NPCs regulate various aspects of the occurrence and progression of liver disease, the cellular heterogeneity and dynamic regulation of NPCs needs to be studied at a single-cell resolution to better understand the pathological mechanism of liver disease.

Single-cell RNA sequencing (scRNA-seq) provides a new perspective for understanding the physiological and pathological processes of multicellular organism (8). By defining the transcriptomic landscape of cells, scRNA-seq can reveal the role of intercellular communication in health and disease at an unprecedented resolution (4). Recently, a diverse range of studies involving scRNA-seq have revealed the heterogeneity of healthy human liver cells (9), explored the distinctive functional composition of infiltrating T cells in hepatocellular carcinoma (10), delineated the transcriptomic landscape and intercellular crosstalk in human intrahepatic cholangiocarcinoma (11), and revealed the heterogeneity of individual cell types and their crosstalk during fibrogenesis in both fibrotic mice and nonalcoholic steatohepatitis (NASH) patients (12). However, a complete single-cell landscape of liver NPCs including health and multiple liver diseases has not been disclosed, and differences in NPCs among these different typical mouse models of liver disease need to be clarified.

In this study, we used the 10x Genomics scRNA-seq platform to profile single cells from healthy mouse and diseased murine livers. The diseased livers were obtained from various mouse models of liver disease, including ALD, 45% high fat-methionine/choline deficient (HF-MCD) diet-induced NASH, bile duct ligation (BDL)-induced cholestatic liver injury, acetaminophen (APAP)-induced DILI, and liver IR injury. Using these data, we aimed to provide a comprehensive transcriptomic overview of NPCs from healthy and diseased murine livers, investigate the heterogeneity of liver NPCs, clarify the differences between the assessed disease models, and developed an interactive website (http://tcm.zju.edu.cn/mlna) to provide universal access to this data source. Together, our findings provide a valuable resource to better understand the pathological mechanisms underlying liver diseases and for clinical therapeutics.

## MATERIALS AND METHODS

### Animals

Eight- to twelve-week-old male C57BL/6J mice were purchased from Charles River Laboratories (Beijing, China). Mice were maintained in specific pathogen-free facilities (12-hour light/dark cycle) with access to food and water *ad libitum*. All animal experiments were performed following procedures approved by the Animal Care and Use Committee of Zhejiang University.

### Animal models of liver disease

#### Model Construction of ALD

The mouse model of ALD by chronic-plus-binge ethanol feeding was described previously (13). Briefly, mice initially received the control Lieber-DeCarli diet (Bio-Serv, Cat#F1259SP) for 5 days to accommodate to a liquid diet, which was followed by acclimation to the ethanol Lieber-DeCarli ethanol liquid diet (Bio-Serv, Cat#F1258SP) of 5% (v/v) ethanol for 2 weeks. On the final day of feeding, an additional gavage of ethanol (5 g kg^-1^, Aladdin Biochemical, Cat#E111993) was administered to mice in the early morning. After 9 hours of gavage, mice were anesthetized for subsequent experiments.

#### Model Construction of NASH

The long-term feeding of choline deficiency combined with high-fat diet was developed to recapitulate key features of human NASH (14). To construct the NASH model, mice were feed on a MCD diet containing 45% kcal fat (Research Diets, Cat#A06071301B) for 8 weeks as previously described (15), which preferable maintained the increase of mice body weight. The normal chow diet fed mice were treated as control.

#### Model Construction of Liver IR Injury

An established mouse model of 70% warm hepatic IR injury was used (16). Briefly, the hepatic artery and portal vein were isolated and clipped with a microvascular clamp, occluding blood supply to the left and middle liver lobes. After 45 minutes of ischemia, the clamp was removed to initiate the reperfusion phase. Mice were sacrificed at 24 hours after reperfusion

#### Model Construction of APAP-induced Acute Liver Injury

Acute liver injury was induced by APAP overdose in mice (17). Before APAP treatment, mice were fasted for approximately 16 hours. Then the animals were intraperitoneally injected once with APAP (300 mg kg^-1^, TCI, Cat#H0190) dissolved in 25% propylene glycol (Sinopharm Chemical Reagent, Cat#30157018) and saline solution. At 24 hours after APAP administration, livers were obtained to be processed for further experiments.

#### Model Construction of BDL-induced Cholestatic Liver Injury

Cholestasis in the experimental model was induced by BDL surgery in mice as previously performed (18). Under general anesthesia, mice were placed supine for midline laparotomy to expose the common bile duct. Then bile duct ligation was performed in two adjacent positions approximately 1 cm from the porta hepatis with 6–0 silk sutures. The duct was then severed by incision between the two sites of ligation. On the tenth day after bile duct ligation, the liver was harvested to isolate NPCs.

### Isolation of Liver NPCs and Preparation of Single-cell Suspensions

Liver NPCs were isolated from mice according to a two-step collagenase method reported previously (6). In detail, murine livers were perfused *in situ* via the inferior vana cava with calcium-free Hank’s Balanced Salt Solution (HBSS, Gibco, Cat#14170112) containing EDTA (0.2 mg mL^-1^, Sigma, Cat#E6758), followed by the buffer II containing pronase (0.4 mg mL^-1^, Sigma, Cat#P5147-1G) and 0.2% collagenase type II (Worthington, Cat#LS004176) at a perfusion rate of 8 mL/minute. Then livers were surgically removed and cut into small pieces. Tissues were transferred in HBSS containing 0.2% collagenase type II, pronase (0.4 mg mL^-1^) and DNase I (0.1 mg mL^-1^, Roche, Cat#10104159001), and then incubated for digestion at 37 °C in a water bath for 20 minutes. DMEM (Mediatech, Cat#10-013-CV) containing 10% serum (FBS, Gibco, Cat#10099-141C) was added at the end of the incubation. Sequentially, hepatocytes removal was achieved by centrifugation for 3 minutes at 50 g. Then cell suspension was filtered using a 40 μm nylon cell strainer (Falcon, Cat#352340). Erythrocytes were lysed by treatment with 3-5 mL ACK lysing buffer (Gibco, Cat#A1049201) for 5 minutes, after which PBS (Beyotime Biotec, Cat#C0221A) was added to terminate the lysis. The resulting suspension was subjected to Dead Cell Removal Kit (Miltenyi Biotec, Cat#130090101) to remove dead cells according to the manufacturer’s recommendations. The obtained cell pellet was washed twice and resuspended in PBS. Cell viability was assessed by Trypan Blue (Gibco, Cat#15250-061).

### 10x Genomics scRNA-seq

Liver NPCs single cell suspensions were loaded onto the 10x Genomics Chromium chip (10x Genomics; Pleasanton, CA, USA) to generate droplets. Then the obtained Gel Beads-in-emulsion (GEMs) were transferred into a PCR tube strip, followed by reverse transcription using ProFlex PCR System (Thermo Fisher, MA, USA). The resulting cDNA was purified and amplified for 12 cycles before cleanup with SPRIselect beads (Beckman, Cat#B23318). Based on the cDNA concentration determined by Qubit (Thermo Fisher, MA, USA), libraries were prepared using the Chromium Single Cell 3’ Library & Gel Bead Kit v3 (10x Genomics; Pleasanton, CA, USA) according to the manufacturer’s instructions. All the libraries were sequenced on the Illumina Novaseq platform by Novogene (Beijing, China).

### Histological Assessment of Liver Sections

Retrieved liver tissues from sacrificed mice were placed immediately in 10% formalin solution. After embedded in paraffin, tissue sections were cut at 5 μm thickness followed by deparaffinization in xylene and rehydration in 100%, 95%, 90%, 80%, 75% alcohol successively. Then the sections were incubated with 3% H_2_O_2_ to inactive endogenous peroxidases for 10 minutes in the dark at room temperature. Nonspecific binding blocking was performed with 5% BSA for 1 hour. Next, slides were stained with hematoxylin and eosin (H&E) for morphological evaluation, and were stained with Sirius Red for fibrosis assessment. For immunochemical analysis, slides were incubated with primary antibodies against LYVE1 (1:2000, Abcam, Cat#218535) at 4 °C overnight in the dark. After three times of PBS washing for 5 minutes, the corresponding *horseradish peroxidase (*HRP)-conjugated Goat anti-Rabbit IgG secondary antibody (Origene, Cat#PV-6002) was sequentially used for incubation at room temperature for 30 minutes. Nuclei were counterstained with hematoxylin. The expression pattern of LYVE1 in liver slides was acquired by Olympus BX63 microscope (Olympus, Shinjuku, Japan) at 200x magnification.

### Immunofluorescence Staining

Liver sections were cut into 5 μm slides from formalin-fixed and paraffin-embedded tissues dissected from C57BL/6J mice. The slides were incubated with the primary antibody against S100A9 (1:500, Abcam, Cat#ab242945) at 4 °C overnight. After slide washing with PBS-T (PBS + 0.05% Tween20), the fluorophore-conjugated secondary antibodies were used for incubation at room temperature for 1.5 hours. For double immunostaining, liver sections were firstly stained with CD31 (1:500, Abcam, Cat#ab182981) or S100A9 (1:500, Abcam, Cat#ab242945) followed by the appropriate secondary antibody. Then LYVE1 (1:1000, Abcam, Cat#ab218535) or YM1 (1:500, Abcam, Cat#192029) antibody was applied onto sections following the identical procedure. Tissue slides were mounted with Antifade Mounting Medium with DAPI (Origene, Cat#ZU9557) for nuclei staining. Fluorescence images were captured with an Olympus BX63 microscope. The fluorophore-conjugated secondary antibodies include Goat anti-Rabbit IgG H&L-Alexa Fluor 488 (1:500, Abcam, Cat#ab150077) and Goat anti-Rabbit IgG H&L-Alexa Fluor 555 (1:500, Abcam, Cat#ab150078)

### Data Processing

The gene expression matrix for each scRNA-seq sample was generated by CellRanger pipeline (10x Genomics) and raw data were processed further in R (version 3.6.1). Quality filtering steps were performed using the Seurat package (version 3.1.2) (19):

1. Genes expressed by less than 5 cells were excluded from further analysis.
2. Cells with fewer than 200 genes expressed, and > 20% of total expression from mitochondrial genes were filtered out.

Through the above steps, 197,194 cells were used for next analysis. For filtered gene expression matrices, gene counts for each cell were normalized by dividing by the total counts for that cell and multiplying by the scale.factor (10,000) with the NormalizeData function of the Seurat package. In order to remove the batch effect, the top 2,000 highly variable genes were used for canonical correlation analysis (CCA) implemented in Seurat.

### Clustering and Cell Typing

After aligning the top 20 dimensions according to CCA, principal component analysis (PCA) was used to reduce dimension using the RunPCA function, and unsupervised clustering was applied using the FindNeighbors function and the FindClusters function with default parameters. Cells were later visualized using the RunTSNE function with the t-distributed stochastic neighbor embedding (t-SNE) algorithm. We then calculated the top marker genes for each cluster using the Wilcoxon rank-sum test, by the FindAllMarkers function (logfc.threshold = 1.5, min.pct = 0.25). The identity for each cluster was annotated based on the SingleR package (version 1.0.5)(20) and the prior knowledge of biology. The cells expressing high levels of classic hepatocyte marker genes (*Alb, Apoa2, Apoc3* and *Mup3*) were filtered out. A total of 171,814 cells remained finally. For sub-clustering of each major liver cell type, a higher “resolution” parameter of FindClusters function was applied. We also used the FindAllMarkers function (logfc.threshold = 0.25, min.pct = 0.25, test.use = “wilcox”) to perform differential expression analysis for each subcluster.

### Pseudo-cell Analysis

As described and confirmed before (21), we performed pseudo-cell analysis to increase the gene expression correlation from high-throughput scRNA-seq data. Briefly, we built a new gene expression matrix for each cell type by constructing pseudo-cells, which were the averages of 20 cells randomly chosen.

### Transcription Factor-target Gene Network Analysis

The regulatory network analysis was performed on pseudo-cell gene expression matrices using the SCENIC package (version 1.1.2-01, corresponds to GENIE3 1.8.0, RcisTarget 1.6.0 and AuCell 1.8.0)(22) with default parameters. Two gene-motif rankings databases of mouse (10 kb around the TSS and 500 bp upstream of TSS) were selected for RcisTarget. To determine the number of “on/off” regulons in each cell type on different models, we set the criteria as follow:

1. we used “mean (AUC scores)” for each regulon as threshold to binarize the regulon activity scores and created the binary regulon activity matrix, where 1 for “on” and 0 for “off”.
2. In each cell type, if the binary activity of the regulon was “on” in more than half of cells, we considered this regulon was “on” in the cell type.

The transcription factor-target gene network was visualized with Cytoscape (version 3.7.2).

### Deconvolution of Liver Microarray/Bulk-seq Data

To accessed the MPs composition in different liver diseases, we applied deconvolution analysis on publicly available microarray/bulk-seq data from annotated liver biopsy specimens taken across the AH (GSE28619), the APAP induced ALF (GSE120652), the BA (GSE159720), the IR injury (GSE151648) and the NAFLD (GSE48452). In particularly, all control samples from 5 datasets were collected together as control group. We only chose the IRI^+^ samples as IR group and the nonalcoholic steatohepatitis samples as NASH group. MPs from our scRNA-seq data were clustered into KCs, MoMFs and DCs, and signature gene expression profiles of these 3 cell types were used to deconvolve the MPs composition of different liver disease samples using CIBERSORTx (23). The composition of MoMFs of different liver disease samples was later associated with the histological features provided by original research paper.

### Pathway Enrichment Analysis

Pathway enrichment analysis was performed using the Gene Ontology (GO) biological process and pathway terms in Metascape (version 3.5)(24) (http://metascape.org) with default parameters, as well as Ingenuity Pathway Analysis (IPA). The results were visualized with the ggplot2 package (version 3.3.0).

### CCI Analysis

Pseudo-cell gene expression matrices were input to predict CCI based on the pseudo-cell gene expression matrices using CellPhoneDB (version 1.1.0) (25). The ligands or receptors which expressed in at least 10% of cells were considered only. For all ligand-receptor pairs, only those with average expression > 0.1 as well as *p*-value < 0.1 were selected for subsequent prediction. To explore immune activation of non-immune cells in different groups, for each type of non-immune cells (endothelial cell, HSC and cholangiocyte), we selected the shared interactions between it and each immune cell type of six major cell types (B cell, MPs, neutrophil, NK, pDC and T cell). We also analyzed the expression levels of immune genes in non-immune cells in different groups. A totally 2484 immune genes in 17 major categories were obtained from ImmPort database (https://www.immport.org). Based on the result of Sirius Red staining, we chose the ligand/receptor genes in two fibrosis-related categories “TGFβ_Family_Member” and “TGFβ_Family_Member_Receptor” to perform the CCI analysis between non-immune cells and immune cells in BDL and HF-MCD groups.

### Metabolic Analysis

We used the method published before to characterize the metabolic heterogeneity in different cell types and groups (26). Totally 1,664 metabolic genes and 85 pathways were obtained from the KEGG database (http://www.kegg.jp), and the metabolic pathways were further grouped into specific categories based on KEGG classifications. The pathway activities were calculated following the protocol using the pseudo-cell gene expression matrices.

### Pseudotime Analysis

Pseudotime analysis was performed on neutrophil (Neu1, Neu2, and Neu3) and kupffer subtypes (MP1, MP2, and MP3) using the monocle R package (version 2.14.0) (27). Genes expressed in less than 10 cells were removed. After reducing the dimensionality of the data using the DDRTree dimensionality reduction algorithm with the reduceDimension function, cell ordering was performed by the orderCells function to build the trajectory. We then calculated the genes differentially expressed along the pseudotime with the differentialGeneTest function and selected those with significant differences (*q*-value < 0.05). For Kupffer cell, we also verified the trajectory and its directionality using the velocyto (version 0.17) (28). We generated annotated spliced and unspliced reads matrices from 10x bam files and selected the aimed cells based on the cell typing result to estimate cell velocities. We set the neighborhood size as 500 cells and all other parameters were default.

### Statistical Analyses

Marker genes for each cell cluster were calculated by the FindAllMarkers function in Seurat R package using the Wilcoxon rank-sum test. Genes with *q*-value < 0.05 were considered statistically enriched in a cluster. For metabolic pathways analysis, we evaluated the activities of metabolic pathways in a specific cell type by using the random permutation test. Only the pathways with *p*-value < 0.05 were considered statistically changed in different cell types or groups. For CCI analysis, the interaction with *p*-value < 0.1, mean expression > 0.1 was indicated statistically significant. For calculating the genes differentially expressed along the trajectory in pseudotime analysis, the genes with significant differences were selected based on *q*-value < 0.05. For plasma biochemical parameters, the significance of differences between control group and model group was determined by two-tailed Student’s *t*-test using Graph-Pad Prism v5.0, the *p*-value < 0.05 was considered statistically significant.

## RESULTS

### An overview of the single-cell atlas of murine liver NPCs

To obtain an overall landscape and compare the cell heterogeneity of liver NPCs in various disease models at a single-cell level, we successfully established 5 classic models of liver disease and performed scRNA-seq on livers from these disease groups and a control group using the 10x Genomics platform (Figure 1A; Supplementary Figure S1A and Table S1). We subsequently developed an interactive website “Murine liver NPCs Atlas” to provide universal access to this data source. Through the browsing function, we can know the expression of clinically important genes related to diseases in different cell types.

**Figure 1.**
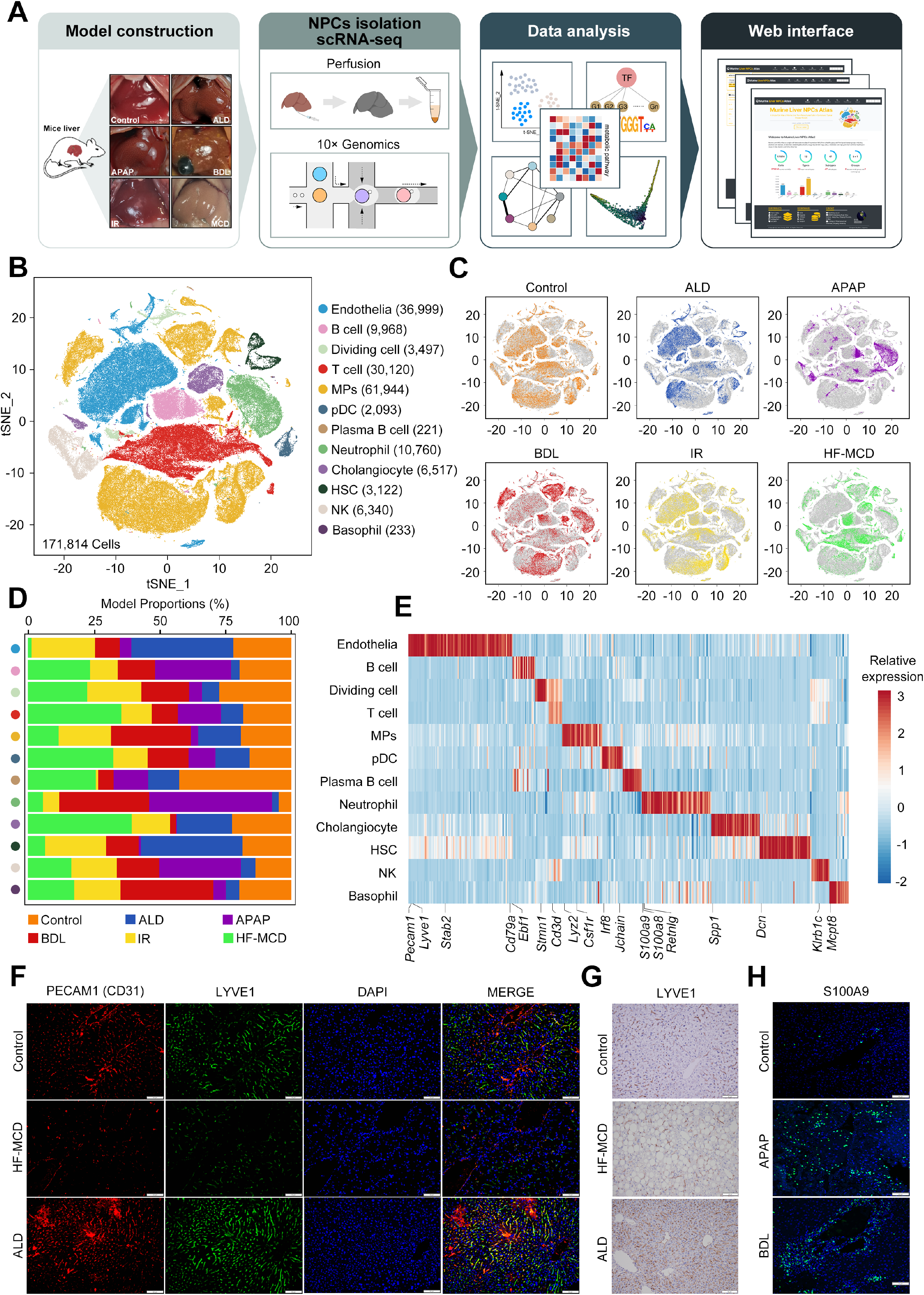
Single cell RNA-seq analysis of murine liver NPCs isolated from different groups. **(A)** Illustration of the study design. **(B)** t-SNE plot visualization of 12 major cell types based on 171,814 single-cell transcriptomes. MPs, mononuclear phagocytes; pDC, plasmacytoid dendritic cell; HSC, hepatic stellate cell; NK, nature killer cell. **(C)** Annotation by different groups. ALD, model of alcoholic liver disease; APAP, model of APAP-induced acute liver injury; BDL, model of bile duct ligation-induced cholestatic liver injury; IR, model of liver ischemia-reperfusion injury; MCD, model of non-alcoholic steatohepatitis. **(D)** Group proportions of the 12 major cell types. **(E)** Gene expression heatmap of the marker genes (logFC > 1.5) for each cell type. **(F)** Immunofluorescence staining of ECs markers (CD31 and LYVE1) in murine livers of control, ALD and MCD groups. Nuclei were stained using DAPI (blue). Scale bars, 50 μm. **(G)** Immunohistochemistry of LYVE1 expression in murine livers of control, ALD and MCD groups. Scale bars, 50 μm. **(H)** Immunofluorescence staining of neutrophil marker S100A9 in murine livers of control, APAP and BDL groups. Nuclei were stained using DAPI (blue). Scale bars, 50 μm.

All liver cells were isolated from mice according to a previously described method, which has a higher collection rate for NPCs (6). After quality control analysis, we obtained 197,194 single-cell transcriptomes in total, which include 25,380 hepatocytes and 171,814 NPCs from 18 mice (3 mice per group) in the control and 5 liver disease groups (Figure 1B and C; Supplementary Figure S1B and C). Clustering analysis of NPCs was subsequently performed using gene expression profiles. The t-distributed stochastic neighbor embedding (t-SNE) plot was used to visualize 12 major clusters of NPCs based on the expression of marker genes (Figure 1B and E; Supplementary Figure S1D). The distribution and proportions of the 12 major cell types in each group provide an insight into the disease groups. For example, an obvious opposite trend was observed in the numbers of ECs in the ALD and HF-MCD groups, and neutrophils were notably increased in the APAP and BDL groups compared with the other groups (Figure 1D). Likewise, the numbers of MPs in the BDL group showed a difference in distribution compared with those in the control group, indicating that some subclusters of MPs may be increased in the BDL group (Figure 1B and C). Clustering analysis of 12 NPCs clusters confirmed several unique transcriptomic characteristics (Figure 1E). For instance, PECAM1 (CD31) and LYVE1 are already well-known ECs markers, and S100A9 is often used as a neutrophil marker (6,29). Consistent with our analysis results, an immunofluorescence staining experiment with PECAM1 and LYVE1 and immunohistochemical staining of LYVE1 confirmed the opposite trend in ALD and HF-MCD groups (Figure 1F and G; Supplementary Figure S1E). Similarly, a dramatic increase in the neutrophil number in the APAP and BDL groups was verified by S100A9 staining (Figure 1H and Supplementary Figure S1F). These results provide an overview of the differences in NPCs in the murine livers among different classic liver diseases.

### Reconstruction and heterogeneity of transcriptional regulatory networks in disease groups

To understand the regulatory networks of transcriptional factors (TFs) in cell types and to determine the differences between groups, we predicted the relevant TFs and binarize the activity scores of TFs. We then counted the number of TFs with activity “on” and that of TFs with activity “off” in each cell type. In most cell types, the number of TFs with activity “on” was increased in disease groups (compared with that in the control group), although there were obvious differences among the disease groups (Figure 2A). The heat map of the activity of TFs revealed that, in each cell type (e.g. ECs or neutrophils), the up-regulated TFs demonstrated an inter-group specificity (Figure 2B and C; Supplementary Figure S2).

**Figure 2.**
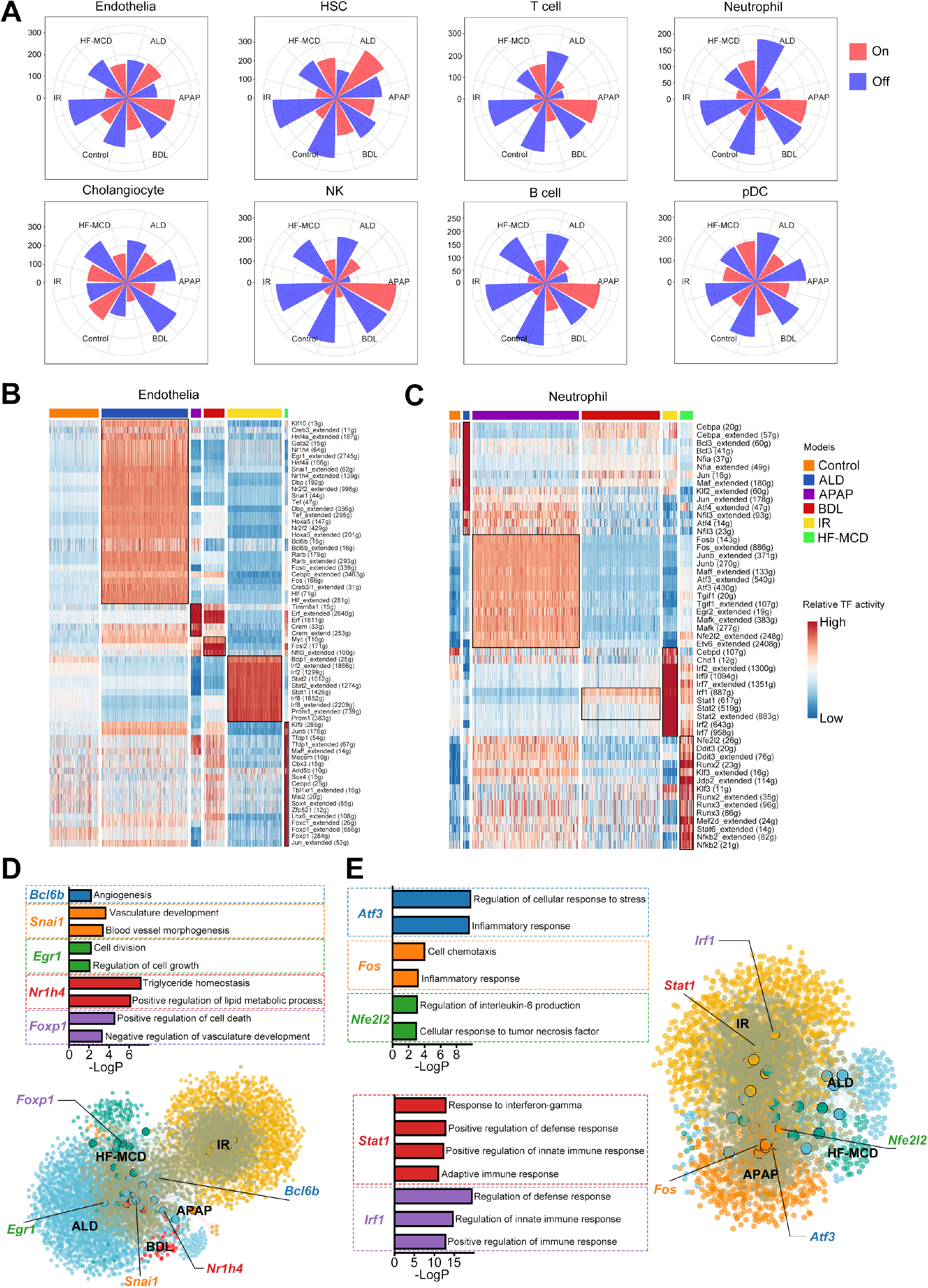
Changes in cellular transcription factor-target gene network in different groups. **(A)** Rose diagrams visualization of the number “on/off” regulons of each cell type in different groups. **(B and C)** Heatmap showing the activity of regulons of ECs **(B)** and neutrophils **(C)** in different groups. Numbers between brackets indicate the potential (extended) target genes for respective TFs. **(D and E)** Network visualization of the inferred transcription factor-target gene networks in ECs **(D)** and neutrophils **(E)**. The octagons represent TFs and the ellipses represent genes. Model-specific TFs showed on **(B)** are represented by different colors. GO analysis of genes regulated by model-specific TFs showing the different functional enrichment.

We found that most TFs for ECs up-regulated in the ALD group were down-regulated in the HF-MCD group observably, including *Egr1, Snai1*, and *Bcl6b*, which are related to angiogenesis, vascular development, and cell growth (Figure 2B and D). However, up-regulated TFs in the HF-MCD group, such as *Foxp1*, are associated with cell death and negative regulation of vascular development according to gene ontology (GO) analysis (Figure 2B and D). These results may partially explain the observed opposite trend of ECs numbers between the ALD and HF-MCD groups presented in Figure 1D. In the IR group, up-regulated TFs (*Irf2, Irf8, Stat1*, and *Stat2*) of ECs are related to the defense response and innate immune response based on the GO analysis (Figure 2B). This is consistent with the characteristics of IR injury. Although inflammation-related neutrophils were dramatically increased in both APAP and BDL groups compared to the control group, neutrophil TFs were differentially regulated (Figure 1D and Figure 2C). TFs (*Fos, Atf3*, and *Nfe2l2*) related to acute inflammation and oxidative stress were up-regulated in the APAP group and down-regulated in the BDL group (Figure 2C and E). In contrast, TFs (*Irf1, Stat1*, and *Stat2*) related to innate immunity, adaptive immunity, and defense response were up-regulated in the BDL group and down-regulated in the APAP group (Figure 2C and E). In the IR group, the TFs up-regulated in neutrophils were similar to those up-regulated in ECs (Figure 2B, C, and E). Together, these data demonstrate that specific diseases reprogram transcriptional regulatory networks and that there is a heterogeneity among disease groups.

### The inter- and intra-group heterogeneity of NPCs

Here, we performed detailed subtype annotation and functional analysis mainly on ECs, HSCs, neutrophils, T cells, MPs, and cholangiocytes. ECs were divided into six subtypes at a higher t-SNE resolution based on their unique transcriptomic signatures: four subtypes (Endo1-Endo4) of LSECs and two subtypes (Endo5 and Endo6) of pericentral (Endo-pc) and periportal (Endo-pp) ECs (Figure 3A-C). LSECs are characterized by two known LSECs markers (*Gpr182* and *Fcgr2b*) (30). Although transcriptomes of LSECs are similar in general, some genes are highly expressed in a subtype-specific manner (Figure 3C). In addition to two conventional LSEC subtypes (Endo1 and Endo2), GO analysis demonstrated that Endo3 and Endo4 are related to the inflammatory response and adaptive immune response, respectively. Furthermore, *C1qa* (a marker of Endo3) was found to stimulate ECs proliferation and promote new vessel formation (31). The group proportions in ECs subtypes were similar to those in ECs (Figure 1D and Figure 3C). Liver fibrosis-associated HSCs were divided into four subtypes, containing quiescent HSCs (HSC1) and activated HSCs (HSC2, HSC3, and HSC4) based on specific markers (Figure 3D-F). Interestingly, GO analysis showed distinguishable functions of HSC2 and HSC3, HSC2 are related to wounding response and ECM organization, while HSC3 are related to inflammatory response and cholesterol metabolic process. All three neutrophil subtypes (Neu1, Neu2, and Neu3) with distinct transcriptomic signatures were dramatically increased in APAP and BDL groups (Figure 3G-I). Evidence is accumulating that neutrophils have different phenotypes and characteristics even in a highly mature state (32). According to the IPA, Neu1 promotes an inflammatory response and cell migration, while Neu2 and Neu3 downregulate inflammation and cell movement (Figure 3J). Mmp8 (a marker of Neu2) is highly expressed in mature neutrophils and to play a beneficial role in chronic and cholestasis liver injury by alleviating fibrosis (33). Likewise, Chil3 (Ym1, a marker of Neu3) is a known marker of M2 macrophages, and Ly6g^*+*^ neutrophils play an anti-inflammatory role in allergic mice (34). Furthermore, our immunofluorescence staining experiment has verified the abundant presence of protective Neu3 subtype in BDL and APAP groups (Figure S4). The subtypes of infiltrated neutrophils in the tissue demonstrates the development of neutrophil heterogeneity and reprogramming of neutrophils from a pro-inflammatory phenotype to an anti-inflammatory phenotype (35). To determine whether neutrophils have polarization for homeostasis maintenance in response to inflammation, we analyzed the pseudotime polarization trajectory of these three neutrophil subtypes. We found that the infiltrated neutrophils demonstrate a polarization trajectory from Neu1 to Neu3 (Figure 3K). Hence, our results demonstrate the existence of protective neutrophil subtypes and neutrophil polarization.

**Figure 3.**
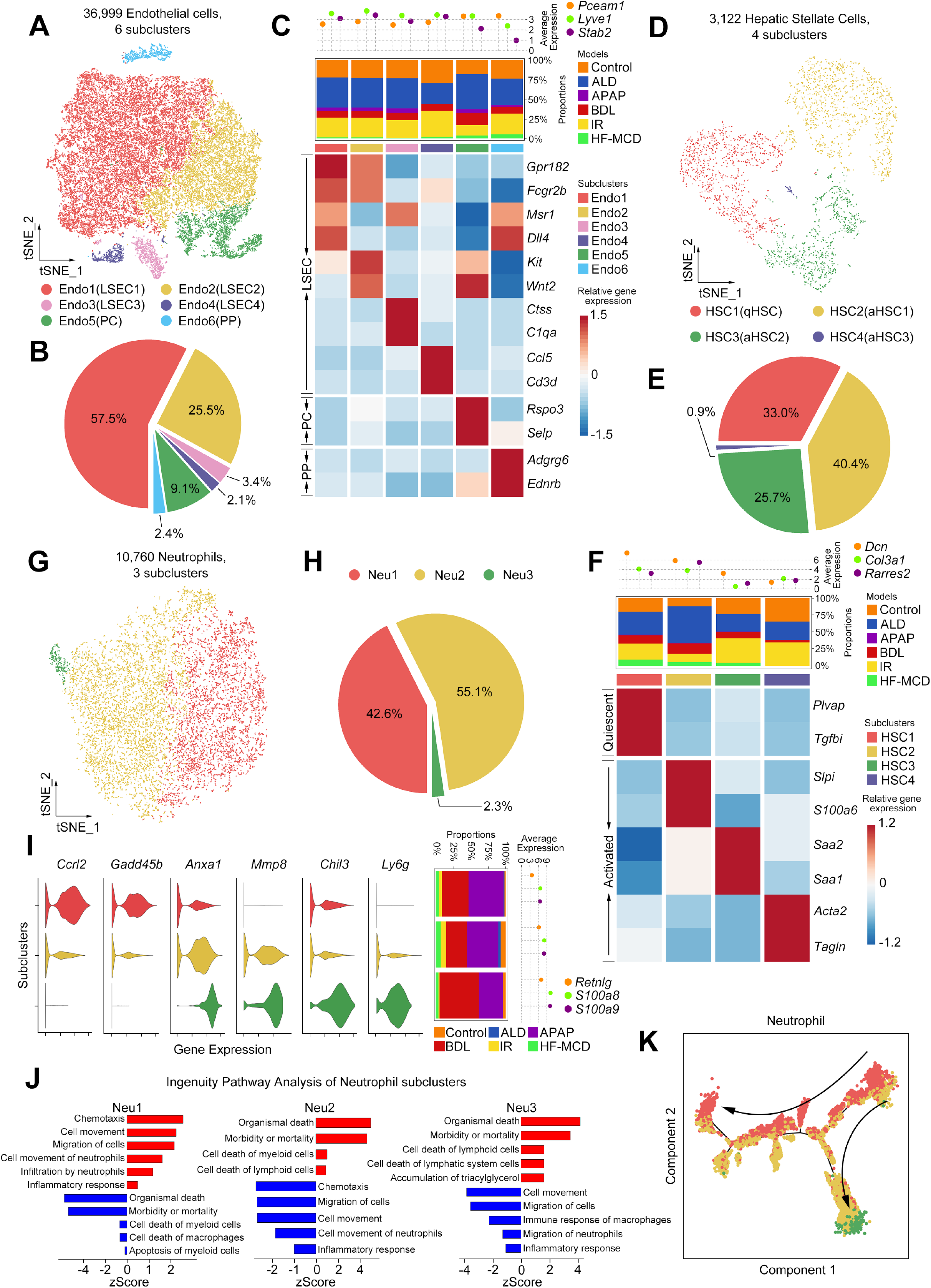
Subcluster analysis of endothelial cell, HSC, and neutrophil. **(A)** t-SNE plot of 36,999 ECs, color-coded by cell subtypes. **(B)** Pie plot showing the proportion of different ECs subtypes. **(C)** Complex heatmap of selected marker genes in each endothelial cell subtype. Top: average expression of known ECs markers; Middle: model proportions of each subtype; Bottom: relative expression of marker genes associated with each cell subtype. LSEC, liver sinusoidal endothelial cell; PC, pericentral endothelial cell; PP, periportal endothelial cell. **(D)** t-SNE plot of 3,122 HSCs, color-coded by cell subtypes. **(E)** Pie plot showing the proportion of different HSC subtypes. **(F)** Complex heatmap of selected marker genes in each HSC subtype. Top: average expression of known HSC markers; Middle: model proportions of each subtype; Bottom: relative expression of marker genes associated with each cell subtype. **(G)** t-SNE plot of 10,760 neutrophils, color-coded by cell subtypes. **(H)** Pie plot showing the proportion of different neutrophil subtypes. **(I)** Complex violin plot of selected marker genes in each neutrophil subtype. Left: expression of marker genes associated with each cell subtype; Middle: model proportions of each subtype; Right: average expression of known neutrophil markers. **(J)** Ingenuity Pathway Analysis of each neutrophil subtype. **(K)** Pseudotime analysis of neutrophils showing the trajectory from N1 to N2.

Similarly, 30,120 T cells were divided into seven subtypes, including natural killer T (NKT) cells (T1), CD4^+^ T cells (T2, T3, and T4), CD8^+^ T cells (T5 and T6), and specific Ramp1^+^ T cells (T7), with distinct transcriptomic signatures (Figure 4A-C). In particular, the number of subtypes T3 (CD4^+^ Foxp3^+^ regulatory T cells) and T6 (effector memory CD8^+^ T cells) was notably elevated in the HF-MCD group, which consistent with previous studies that activated CD4^+^ and CD8^+^ T cells is essential for the progression of NASH and liver fibrosis (Figure 4C) (14,36). T4 and T5 are naive CD4^+^and CD8^+^ T cells with high expression of *Ccr7* and *Sell* (10). Finally, a specific subtype of Ramp^+^ T cells was also identified, which plays a role in angiogenesis according to GO analysis. The role of T cells in angiogenesis and vasculogenesis has previously been noted under pathological and physiological conditions (37). MPs, the largest cell type of NPCs, were divided into eight subtypes based on their unique transcriptomic markers, including KCs (MP1-MP4), MoMFs (MP5 and MP6), and conventional dendritic cells (cDCs) (MP7 and MP8) (Figure 4D-F) (4,6,38). MP3 is a type of periodic KC characterized by high expression of *Stmn1* (Figure 4F) (38). Trem2 and Chil3 are well-known markers of pro- and anti-inflammatory macrophages, respectively (6,38). Ingenuity Pathway Analysis (IPA) analysis was used to confirm that *Trem2*^+^ MoMFs (MP5) up-regulated the inflammatory response pathways and *Chil3^+^* MoMFs (MP6) down-regulated those inflammatory response pathways (Figure 4G). Cholangiocytes are an important type of intrahepatic NPCs that participate in bile production and homeostasis (39). Cholangiocytes were divided into four subtypes (Cho1-Cho4) using distinctive transcriptomic markers (Figure 4H-J). Besides, cell types including NK cells (Figure S3A-C), dividing cells (Figure S3D-F), B cells (Figure S3G-I), and plasmacytoid dendritic cells (pDCs) (Figure S3J-L) were divided into different cell subtypes according to their individual transcriptomic signatures. Altogether, above results demonstrate the heterogeneity of NPCs in the liver.

**Figure 4.**
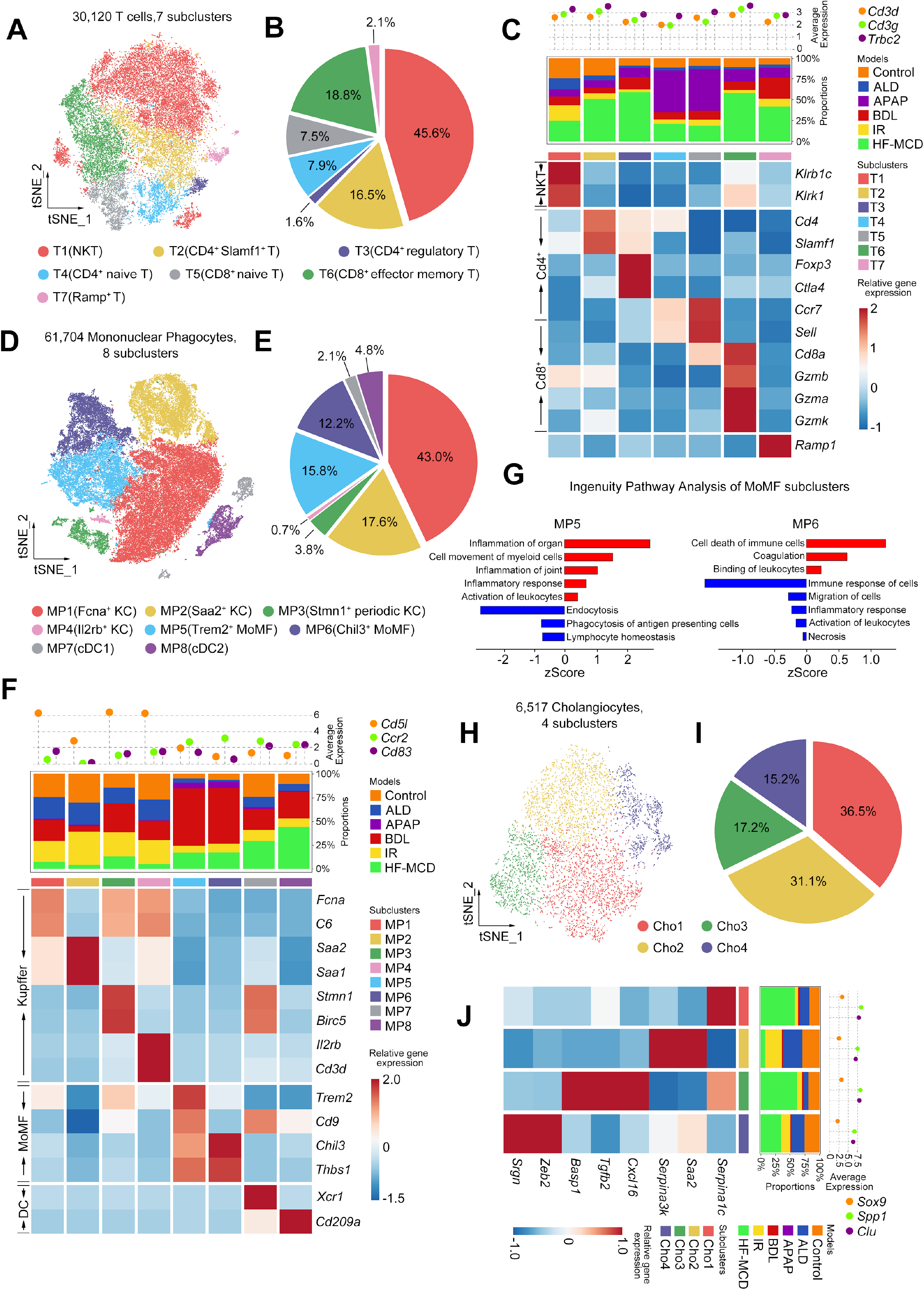
Subcluster analysis of T cell, MPs, and cholangiocyte. **(A)** t-SNE plot of 30,120 T cells, color-coded by cell subtypes. **(B)** Pie plot showing the proportion of different T cell subtypes. **(C)** Complex heatmap of selected marker genes in each T cell subtype. Top: average expression of known T cell markers; Middle: model proportions of each subtype; Bottom: relative expression of marker genes associated with each cell subtype. NKT, nature killer T cell; Cd4^+^, Cd4^+^ T cell; Cd8^+^, Cd8^+^ T cell. **(D)** t-SNE plot of 61,704 MPs, color-coded by cell subtypes. **(E)** Pie plot showing the proportion of different MPs subtypes. **(F)** Complex heatmap of selected marker genes in each MPs subtype. Top: average expression of known MPs markers; Middle: model proportions of each subtype; Bottom: relative expression of marker genes associated with each cell subtype. Kupffer, kupffer cell; MoMF, recruited monocyte-derived macrophage; DC, dendritic cell. **(G)** Ingenuity Pathway Analysis of each MoMF subtype. **(H)** t-SNE plot of 6,517 cholangiocytes, color-coded by cell subtypes. **(I)** Pie plot showing the proportion of different cholangiocyte subtypes. **(J)** Complex heatmap of selected marker genes in each cholangiocyte subtype. Left: relative expression of marker genes associated with each cell subtype; Middle: model proportions of each subtype; Right: average expression of known cholangiocyte markers.

As recruited macrophages, MoMFs play important regulatory roles in a variety of liver injury (3). To determine whether MoMFs expand in various human liver diseases as we found in mouse models, we analyzed liver RNA sequencing data of patients with NASH, alcoholic hepatitis (AH), IR injury after liver transplantation, APAP-induced acute liver failure (ALF), and biliary atresia (BA). We applied differential gene expression signatures of KCs, MoMFs, and DCs to the deconvolution algorithm to evaluate the composition of MPs in human liver (Supplementary Figure S5A and B). Results showed abundant expansion of MoMFs in patients with NASH, IR injury, APAP-induced ALF, and BA, which is consistent with our findings (Supplementary Figure S5A-C). With the increase of MoMFs, the histological NAFLD activity score (NAS), fibrosis score, and inflammation score deteriorated (Supplementary Figure S5D). It indicated that the expansion of MoMFs positively correlated with the progress of NASH and the degree of fibrosis.

Furthermore, one of the advantages of scRNA-seq technology is to understand the characterization of gene expression in different cell types/subtypes. In order to determine the expression characteristics of disease-related clinically significant genes in different cell types/subtypes, we analyzed the gene expression of a series of blood markers and drug targets for NASH diagnosis and therapy in each cell type. Cytokeratin 18 (CK18) is a blood marker of apoptosis and fibrosis for diagnosis of NASH (40). We found that *Krt18*, which encodes CK18, is specifically high expressed in cholangiocytes and obviously increased in subtype Cho3 in HF-MCD group (Supplementary Figure S5E and F), suggesting that NASH is accompanied by significant injury and apoptosis of cholangiocytes, especially subtype Cho3, and cholangiocytes may associate with the mechanism of NASH. Galectin 3 (Gal-3) is one of the promising targets involved in fibrosis for NASH treatment in clinical trials (40). We found that the gene expression of *Lgals3* (encoding Gal-3) is increased in cell types of MPs and HSCs in the HF-MCD group (Supplementary Figure S5G). Moreover, the expression of *Lgals3* is elevated in subtypes of KCs, MoMFs, and HSC2 in the HF-MCD group (Supplementary Figure S5H), implying that it not only reminds the importance of these three cell subtypes to liver fibrosis, but also provides more precise guiding significance for the development of drugs with Gal3 as the drug target. Altogether, these indicated the clinical value of our scRNA-seq data.

### Metabolic reprogramming of NPCs across disease groups

Considering that the liver plays an important role in regulating energy metabolism in the whole body, the metabolic reprogramming of NPCs in liver disease states is worth investigating. The hepatic immune response includes the enhanced glucose metabolism of immunocompetent cells (41). HSCs show a particularly high sensitivity, and they play an important role in immune metabolism by maintaining liver function and responding to injury (42). Next, we investigated the features of metabolic pathway reprogramming of NPCs in different disease groups by quantifying metabolic pathway activity based on a previously described pathway activity score (26). The pathways investigated included those for carbohydrate metabolism, energy metabolism, lipid metabolism, etc. Almost all metabolic pathways in cholangiocytes were dramatically activated in the BDL group (Figure 5A). This is consistent with the pathological mechanism of cholestatic liver injury induced by ligation of the bile duct, in which the siltation and reflux of bile aggravates cholangiocyte stimulation and subsequent damage. In addition, numerous metabolic pathways in HSCs and MPs, including the carbohydrate, energy, lipid, etc. metabolic pathways, were activated (compared with control) in the BDL, IR, and HF-MCD groups (Figure 5A and B). This suggests that the metabolic activation of HSCs and MPs is required to exert an appropriate immune effect. Interestingly, NK cells were only activated in the IR group, suggesting that NK cells have specific metabolic activity in IR liver injury (Figure 5B). Next, through analysis of the metabolism in subtypes of MPs, HSCs, and NK cells in each disease group, we found that the metabolic activity of subtypes KCs, HSC2, and NK3 was the strongest in BDL, IR, and HF-MCD groups (Supplementary Figure S6A-C).

**Figure 5.**
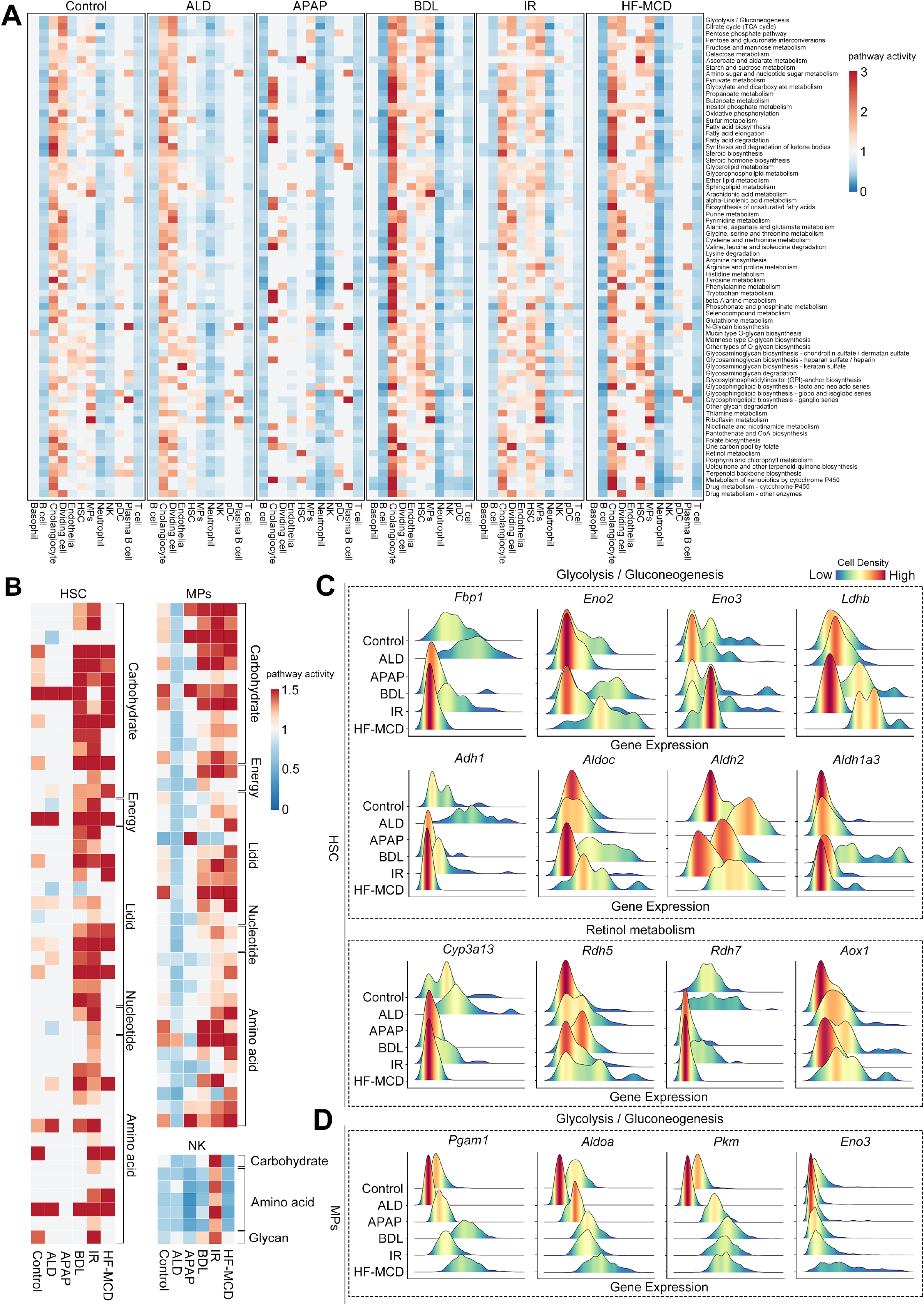
Disease-specific metabolic reprogramming of each cell type. **(A)** Metabolic pathway activities of each cell type in different groups. For each metabolic pathway, the pathway activity scores larger than 1 or smaller than 1 means significantly upregulated or downregulated. **(B)** Metabolic pathway activities in HSC (left), MPs (top right) and NK (bottom right) in different groups. **(C)** Mountain map visualization of the expression of glycolysis/gluconeogenesis pathway related genes (top) and retinol metabolism pathway related genes (bottom) in HSC in different groups. Color-coded by cell density. **(D)** Mountain map visualization of the expression of glycolysis/gluconeogenesis pathway related genes in MPs in different groups. Color-coded by cell density.

Energy metabolism is essential in activated HSCs to support a multitude of functions, including proliferation, secretion of ECM and cytokines, and migration to the injury regions. In addition, carbohydrate and lipid metabolism are required for the activation of HSCs, because transdifferentiation into the myofibroblast phenotype requires upregulation of glycolysis and depletion of retinol-containing cytoplasmic lipid droplets to meet energy demands (42). To understand further, we analyzed gene expression of glycolytic and retinol metabolism pathway in each cell type in different groups (Supplementary Figure S7A and B). We observed that the gene expression of inhibition of glycolysis (*Fbp1*) was down-regulated, while genes expression of promotion of glycolysis and retinol metabolism (*Eno2, Eno3* and *Rdh5, Aox1*) were elevated in HSCs in HF-MCD and BDL groups (Figure 5C) (43–45). These results confirm the abovementioned metabolic up-regulation and demonstrate the inter-group heterogeneity of HSCs activation. Besides, the expression of glycolysis-related genes was up-regulated in MPs (to varying degrees) in BDL, IR, and HF-MCD groups (Figure 5D). The expression of these genes in different cell subtypes can be further explored on our website. These findings reveal the heterogeneity of the metabolic reprogramming of NPCs in different liver diseases, especially confirm the importance of metabolic activation of HSCs and MPs in liver disease.

### Inter-group heterogeneity in communication between non-immune cells and immune cells

Cell-cell interaction (CCI) is a basic feature of multicellular organisms, playing an essential role in numerous biological processes (46). The construction of a CCI network based on ligand-receptor interaction is a common strategy for analyzing scRNA-seq data (46). Non-immune NPCs, including ECs, cholangiocytes, and HSCs, also play a role in immune activation by communicating with immune cells and thus influencing pathological progression (47). To evaluate the global participation of non-immune cells in the immune response, we investigated the expression of immune-related genes in ECs, cholangiocytes, and HSCs and constructed a model of the CCI network between non-immune and immune cells in different disease groups (Figure 6). We found that expression levels of immune-related genes, especially those encoding chemokines/cytokines and their receptors, were up-regulated in ECs, cholangiocytes, and HSCs and that the different disease states show differential expression (Figure 6A).

**Figure 6.**
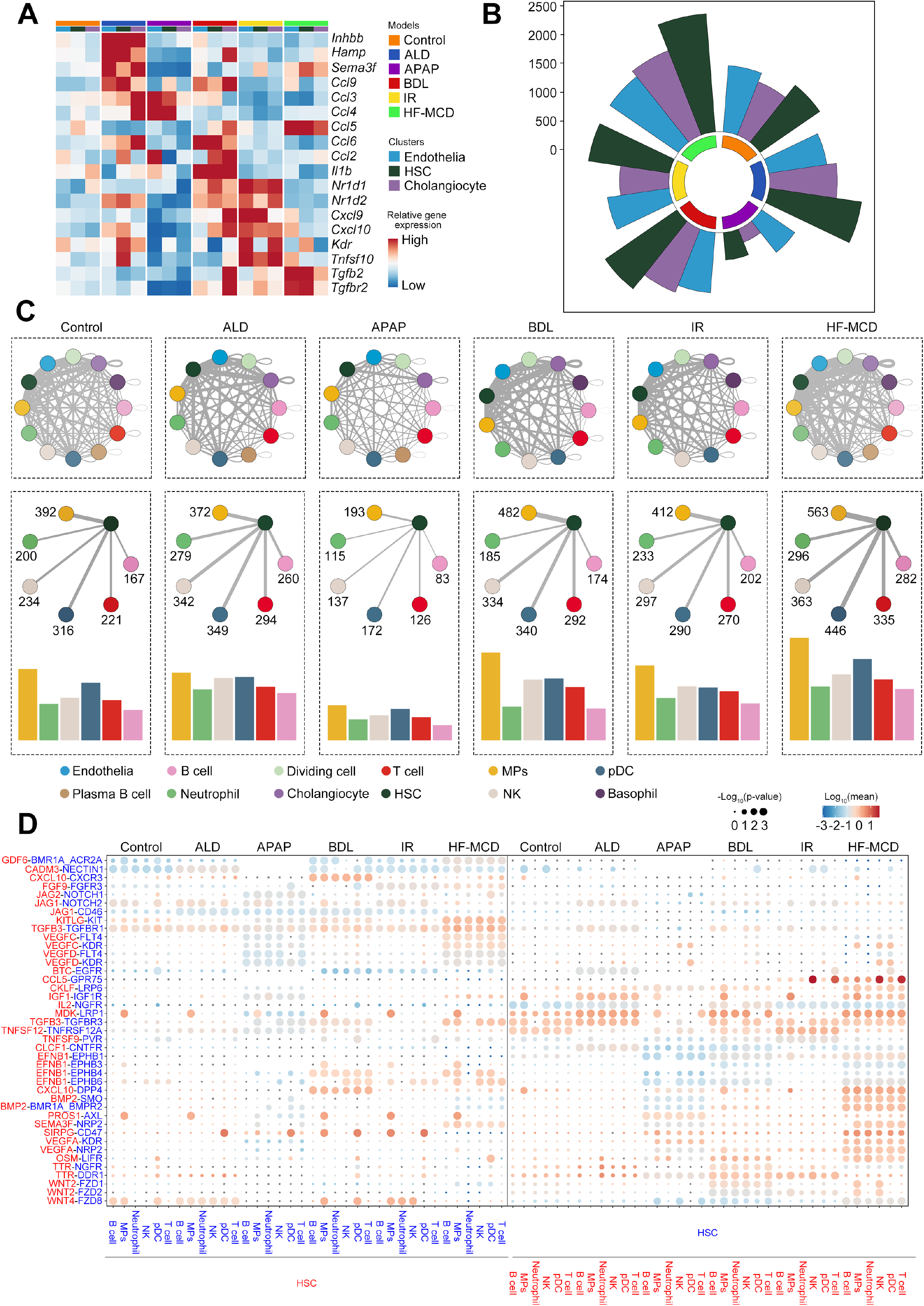
The CCI network between non-immune cells and immune cells in murine livers. **(A)** Heatmap showing the relative expression of immune genes in non-immune cells (endothelial cell, HSC, cholangiocyte) in different groups. **(B)** The number of interaction pairs between non-immune cells and other six immune cells (B cell, MPs, neutrophil, NK, pDC and T cell) in different groups. **(C)** An overview of interaction network between different cells (top) and the interactions between HSC and immune cells (bottom). The line thickness is proportional to the number of interactions between two cell types. **(D)** Dot plot displaying the specific ligand-receptor interactions between HSC and immune cells in different groups. Size of the dot represents statistical significance of the indicated interactions and color of the dot represents the total mean of the individual partner average expression values in the corresponding interacting pairs of cell types.

After comparison of CCI networks between non-immune and immune cells in different groups, we observed a notable increase of CCIs in most disease groups compared to the control group and the number of interactions between HSCs, ECs, cholangiocytes and different intrahepatic immune cells showed an inter-group heterogeneity (Figure 6B and C; Supplementary Figure S8A and B). Moreover, we identified unique ligand-receptor pairs of the CCI in each model group compared with control and found differences in ligand-receptor pairs between model groups (Figure 6D and Supplementary Figure S8C). Functions of unique ligand-receptor pairs in HSCs are mostly related to immunity, inflammatory response, cell proliferation, apoptosis, and transdifferentiation (Figure 6D). Surprisingly, expression level of the ligand-receptor pair CCL5-GPR75 was specifically enhanced, the interaction between them was only observed between immune cells (especially NK and T cells) and HSCs in IR and HF-MCD groups (Figure 6D). These findings indicate the inter-group heterogeneity in the immune activation of non-immune cells.

### Inter-group cell heterogeneity in transcriptional dynamics

KCs are resident macrophages found throughout the mammalian liver and play essential roles in liver disease (47). To investigate the inter-group heterogeneity of KCs polarization process and of KCs transcriptional dynamics, we analyzed polarization trajectories of three KC subtypes using monocle2 and RNA velocity methods for pseudotime ordering. We observed a polarization trajectory from periodic KC subtype MP3 to MP1 to MP2 in all groups (except the APAP group, which had rarely population of KCs) (Figure 7A and B). A similar polarization trajectory was inferred using the RNA velocity method, with KCs polarization in the HF-MCD group being the most obvious (Figure 7B). Next, we investigated genes with dramatically perturbed expression along this trajectory in each group and identified 501 genes common to all groups (Figure 7C and D). Interestingly, we found that the expression of apoptosis-related genes *Bax* and *Bcl2a1b* was up-regulated and of anti-proliferation factors *Btg1* and *Btg2* was markedly reduced along the trajectory in the control group, while completely opposite trend was observed in HF-MCD, BDL, and APAP groups (Figure 7E). Furthermore, as the trajectory changes, expression levels of inflammation-related genes (*Ccl5, Ccrl2, Cxcl2*, and *Trem2*) and fibrosis-related genes (*Tgfb1* and *Tgfbi*) were markedly increased in HF-MCD, BDL, and APAP groups (Figure 7E). Together, these results indicate that the polarization of KCs in the healthy liver is highly correlated with periodic proliferation and apoptosis, while in disease, periodic KCs are polarized into functional KCs to play important roles in progression of liver disease.

**Figure 7.**
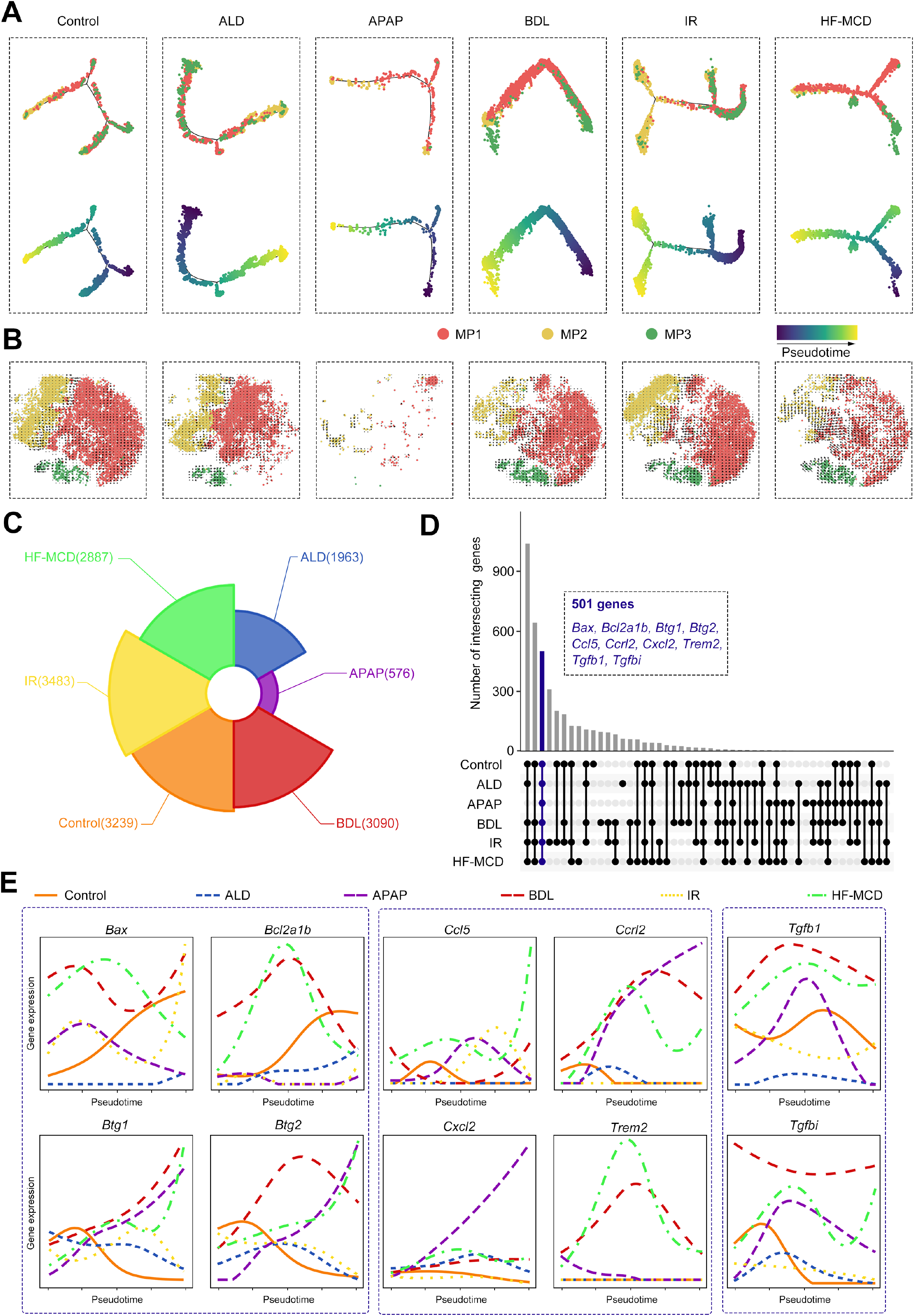
Trajectory analysis of KCs in different groups. **(A and B)** Trajectory inference of 3 KC subtypes (MP1, MP2 and MP3) using monocle **(A)** and RNA velocity **(B)**, colored by cell types or pseudotime. RNA velocity field (black arrows) were visualized on t-SNE plot of 3 KC subtypes. **(C)** The number of genes significantly differentially expressed along the pseudotime in different groups (*q*-value < 0.05). **(D**) Upset plot of intersections between the genes showed in **(C)** in each group. Blue bar: 501 intersecting genes in 6 groups, some of which relate to apoptosis, inflammation and fibrosis were list in blue. **(E)** Expression profiles of apoptosis-related genes (left), inflammation-related genes (middle) and fibrosis-related genes (right) along the pseudotime in 6 groups.

## DISCUSSION

Liver diseases, including DILI, cholestatic liver injury, liver IR injury, ALD, and NASH, are associated with extremely high morbidity and mortality worldwide, causing a huge social burden (2). APAP-induced DILI is the most common and clinically relevant model of intrinsic DILI (48). BDL is the most widely used classic experimental model of cholestasis (49). Liver IR injury has been considered as a potential mechanism responsible for organ dysfunction and injury after liver surgery such as liver transplantation (50). The ALD model was constructed based on a method published in *Nature Protocols*, which describes the generation of a simple and effective ALD model with no mortality rate, no liver fibrosis, marked elevation of alanine aminotransferase and steatosis (13). The pathology of NASH can be induced by the MCD diet rather than a high-fat diet (HFD) in C57BL/6J mice (14). MCD diet is a valuable tool for investigating the inflammatory effects in NASH due to its availability and simplification (51). Inadequate intake of methionine/choline can lead to defective lipoprotein secretion and oxidative stress caused by impaired β-oxidation in the liver, and further induce hepatic steatosis, inflammation and fibrosis (14,51). However, mice fed the MCD diet will not develop any metabolic diseases associated with obesity or insulin resistance, and even loss weight (52). Thus, the MCD diet cannot fully recapitulate the characteristics of NASH patients. Nonetheless, since at least 90% of Americans do not meet the recommended choline intake, and choline deficiency in NASH patients can lead to more severe fibrosis, we applied a 45% HF-MCD diet to investigate characteristics of NASH (53,54).

This study provided new insights into liver physiology and pathology through single-cell transcriptomic technologies. Here, we obtained a comprehensive single-cell transcriptomic landscape of NPCs from livers of healthy and diseased mice, and constructed a website to provide simple access to all our data. Through analysis of distribution and proportions of ECs cluster in each group, we observed that the number of ECs was markedly reduced only in the HF-MCD group, even though both ALD and HF-MCD involve steatosis (Figure 1D, F, and G; Supplementary Figure S1A and E). Thus, ECs injury is a characteristic of the HF-MCD group, which is in agreement with a previous study (6), and ECs injury may related to the degree of liver steatosis and fibrosis. Furthermore, although ECs were greatly reduced in the HF-MCD group, the CCI between ECs and immune cells was notably increased (Figure 6B and Supplementary Figure S8A), indicating that ECs injury is associated with an enhancement in CCI. ECs injury in the HF-MCD group also showed regional heterogeneity, which represented that the damage to LSECs was greater than the damage to Endo-pc and Endo-pp, and the LSEC population was decreased compared with the Endo-pc and Endo-pp populations (Figure 3C).

Neutrophils are considered as main mediators of the inflammatory response during tissue injury. Even though a dramatic infiltration of neutrophils was observed in both APAP and BDL groups, the number of MoMFs was only markedly increased in the BDL group (extremely low in the APAP group), implying that the main cell type involved in the inflammatory response is different in APAP- and BDL-induced liver injury (Figure 3I and Figure 4F). Recent evidence suggests that infiltrated neutrophils can polarize into a protective phenotype, which exerts an anti-inflammatory effect and restores homeostasis (55). Here, we verified the existence of protective neutrophil subtype and analyzed the polarization trajectory of neutrophils from Neu1 to Neu3 subtype (Figure 3K and Supplementary Figure S4). The number of activated CD4^+^ and CD8^+^ T cells was greatly increased in the HF-MCD group (Figure 4C), which is consistent with clinical results suggesting that CD8^+^ T cells are increased in the livers of NASH patients (56). Moreover, NKT cells play a role in the fibrotic progression of NASH, and activation of CD8^+^ T cells and NKT cells can lead to NASH via crosstalk with hepatocytes (14), indicating that T cells play an important role in progressive NASH. In order to further explore the clinical application value of our data, we compared our data with publicly available bulk RNA-seq data from human liver diseases. Due to the limitations of analytical method, we only obtained the consistent result of the increase in the proportion of MoMFs at a lower resolution (Supplementary Figure S5A-C). We can further analyze to complement the present database, when human single-cell data of these liver diseases are available later.

The CCI analysis demonstrated that immune activation of non-immune cells had heterogeneity between different disease groups. In comparison with the control, differences in specific ligand-receptor pairs involved in immune and non-immune cell interactions were observed in each model group. For instance, the interactions between CXCL10 and CXCR3, and CXCL10 and DPP4 (from HSCs to immune cells) were specific for the BDL group, while the interaction between CXCL10 and DPP4 (from immune cells to HSCs) was specific to the HF-MCD group (Figure 6D). *Tgfb2* (transforming growth factor beta 2) is known as a positive regulator of liver fibrosis and is a participant in biliary-induced liver disease based on previous data obtained from a BDL mouse model (57). In agreement with these observations, our results demonstrate that only cholangiocytes in the BDL group showed a high expression of *Tgfb2* and its receptor *TgfbR2*. However, in the HF-MCD group, ECs and HSCs (rather than cholangiocytes) showed a high expression of *Tgfb2* and *TgfbR2* (Figure 6A). Considering that liver fibrosis was obvious in both the BDL and HF-MCD groups, we investigated differences in the gene expression of *Tgfb* family members between these groups (Supplementary Figure S8D and E). *Tgfb* family genes were highly expressed in cholangiocytes in the BDL group, and in HSCs and ECs in the HF-MCD group. Moreover, while fibrosis-related CCIs between cholangiocytes and immune cells in the BDL group were stronger (compared with the HF-MCD group), CCIs between HSCs and immune cells were more obvious in the HF-MCD group (Figure S8F). These results indicate that cholangiocytes are mainly responsible for cholestatic liver fibrosis (58), while HSCs mainly contribute to the fibrosis in NASH. Although the total number of cells collected in the APAP group was the lowest, the reasons for the greatly reduced CCIs in the APAP group need to be explored further.

In conclusion, we first have provided here, a comprehensive single-cell transcriptomic landscape of murine liver NPCs in health and 5 liver disease models (representing more than 70% incidence of liver disease). Although more disease models and the single-cell spatiotemporal heterogeneity of intrahepatic cells, including hepatocytes, should be considered in the future study, this study has prominently increased our understanding of the physiological and pathological mechanisms underlying liver function and dysfunction, and should contribute to the clinical diagnosis and therapeutics of liver diseases.

## DATA AVAILABILITY

All raw single-cell RNA sequencing data in this paper have been deposited into the Gene Expression Omnibus (GEO) database (GEO: GSE166178). The raw or processed data can be downloaded on the GEO database or our website (http://tcm.zju.edu.cn/mlna). Custom code for analysis will be available by request.

## SUPPLEMENTARY DATA

Supplementary Data are available at NAR Online.

## ACKNOWLEDGEMENTS

Z.W. designed, performed, and analyzed all experiments; J.Q. processed scRNA-seq data and performed computational analysis; P.Z., R.G., H.L., and S.Z. participated in the experiment; Z.W., J.Q., P.Z., and J.Y. wrote the manuscript; X.L. and X.F. supported and supervised the experiment and revised the manuscript; X.F. conceptualized the study. All the authors reviewed the manuscript.

