## Supplementary data for "A single-cell transcriptomic atlas characterizes liver non-parenchymal cells in healthy and diseased mice"

### Supplemental Figure S1

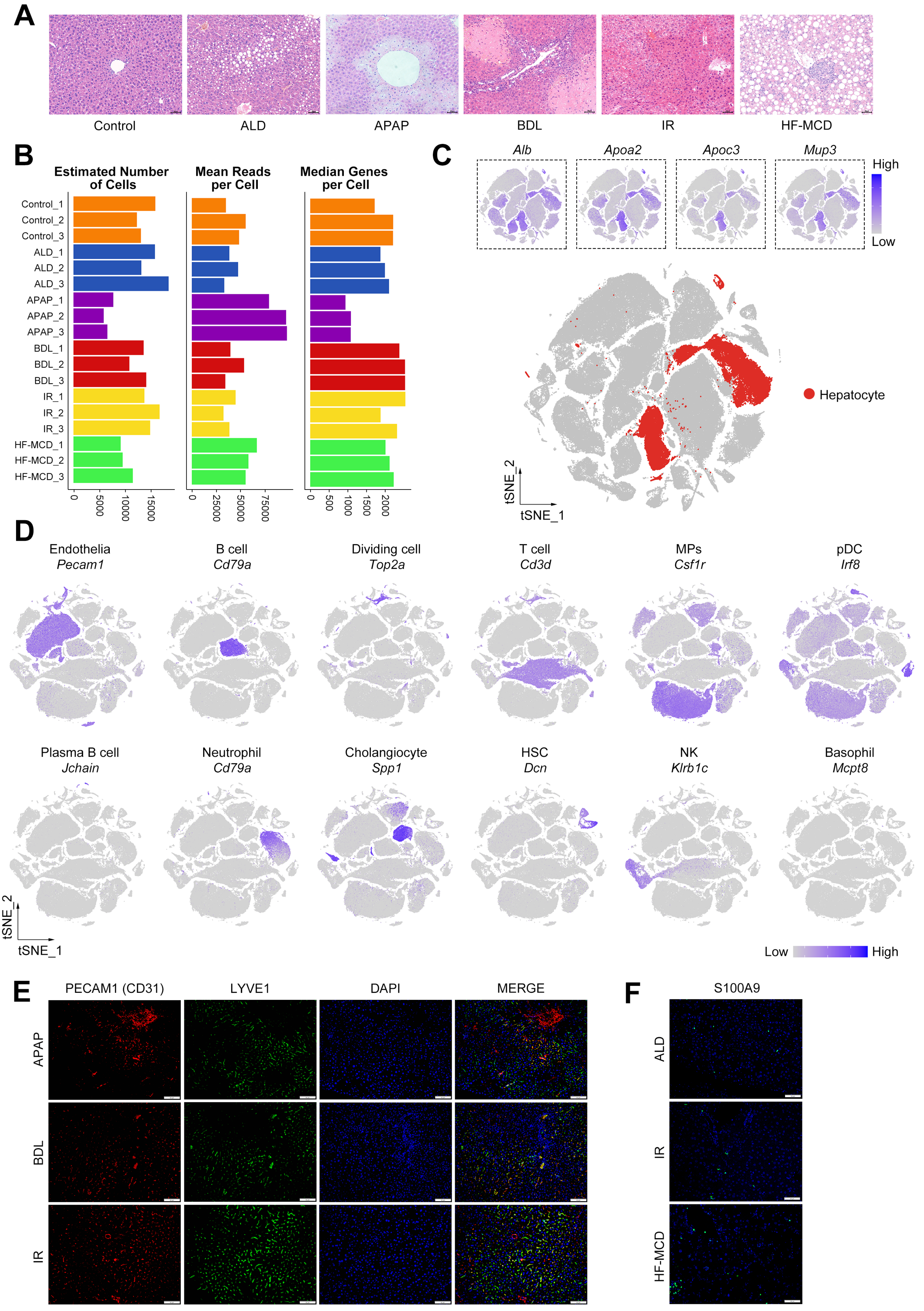

**Supplemental Figure S1.** Single cell RNA-seq analysis of murine liver NPCs isolated from different groups. A) H&E staining morphology of murine livers in 6 groups. Scale bars, 50 μm. B) Estimated number of cells, mean reads per cell and median genes per cell for the scRNA-seq data in the 18 samples across 6 groups. C) t-SNE plot visualization of unsupervised clustering based on 197,194 single-cell transcriptomes. Cells with expressing high levels of classic hepatocyte marker genes (*Alb*, *Apoa2*, *Apoc3* and *Mup3*) are highlighted by red color. D) Feature plot showing the expression of selected marker genes of each cell type. E,F) Immunofluorescence staining of ECs markers (CD31 and LYVE1) (E), and neutrophil marker S100A9 (F) in murine livers of other groups. Nuclei were stained using DAPI (blue). Scale bars, 50 μm. Related to figure 1.

### Supplemental Figure S2

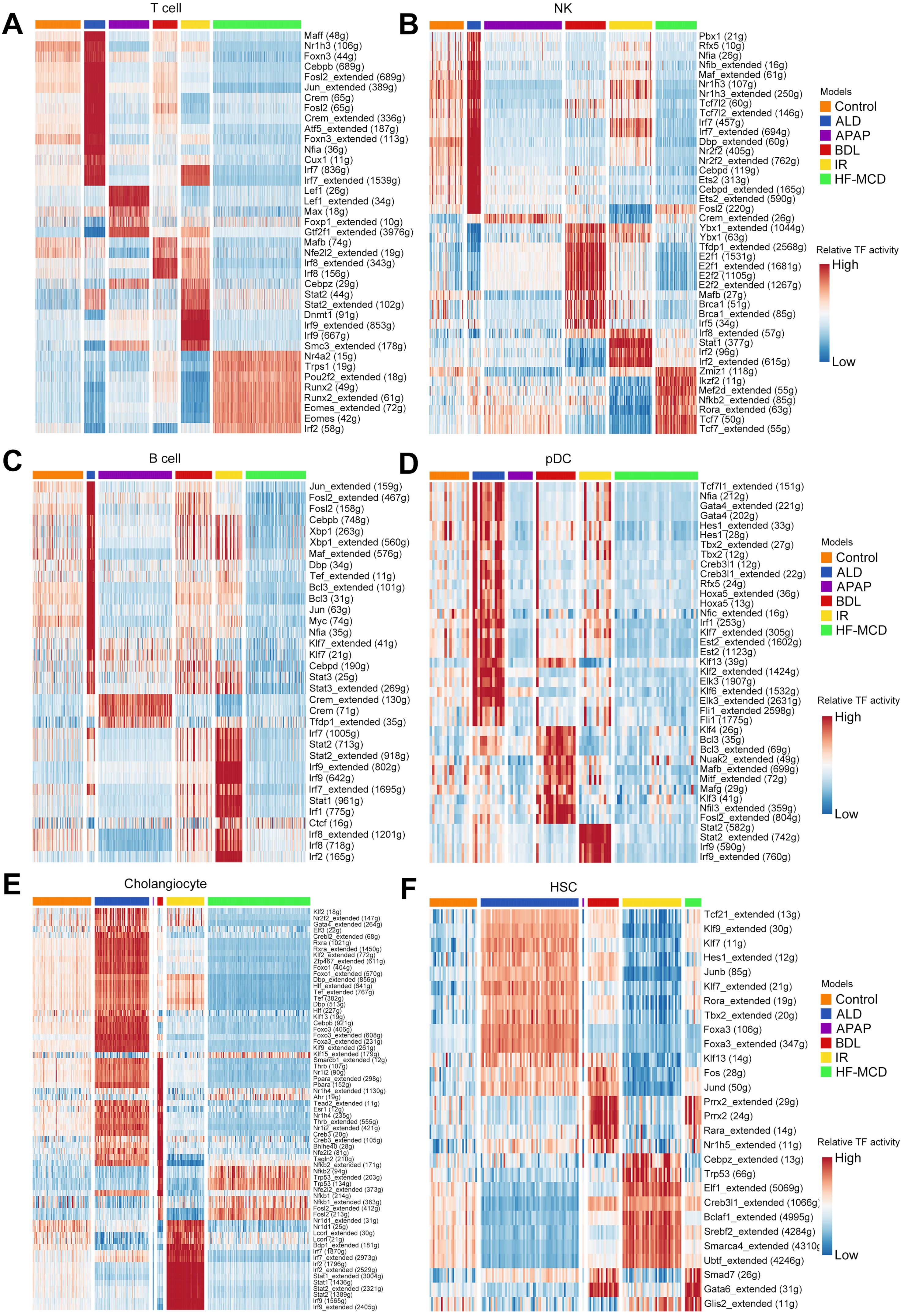

**Supplemental Figure S2.** Changes in cellular transcription factor-target gene network in different groups. Heatmap showing the activity of regulons in T cell (A), NK (B), B cell (C), plasma B cell (D), cholangiocyte (E) and HSC (F) in different groups inferred by SCENIC. Numbers between brackets indicate the potential (extended) target genes for respective TFs. Related to figure 2.

### Supplemental Figure S3

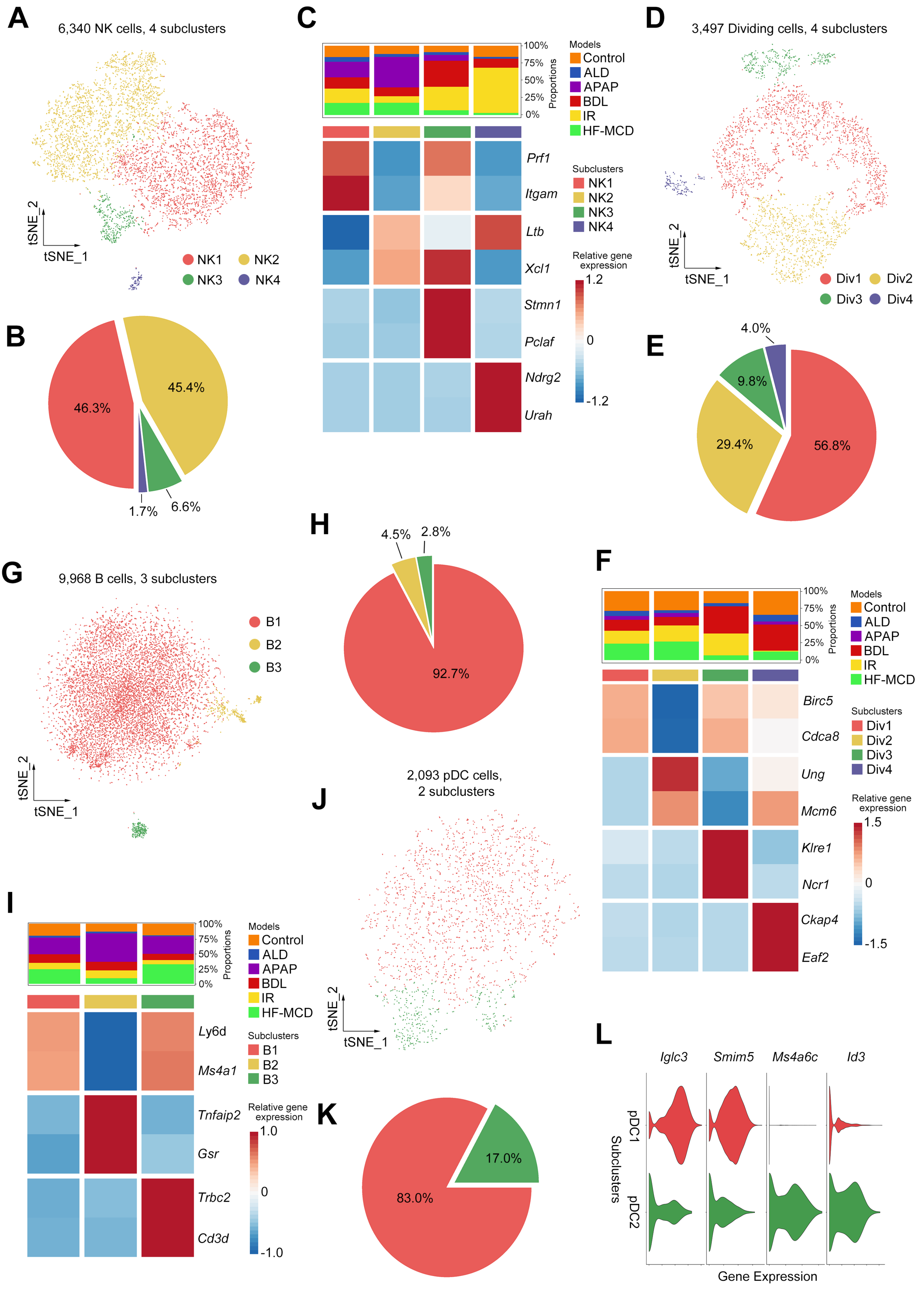

**Supplemental Figure S3.** Subcluster analysis of NK, dividing cell, B cell and pDC. A) t-SNE plot of 6,340 NK cells, color-coded by cell subtypes. B) Pie plot showing the proportion of different NK subtypes. C) Complex heatmap of selected marker genes in each NK subtype. Top: model proportions of each subtype; Bottom: relative expression of marker genes associated with each cell subtype. D) t-SNE plot of 3,497 dividing cells, color-coded by cell subtypes. E) Pie plot showing the proportion of different dividing cell subtypes. F) Complex heatmap of selected marker genes in each dividing cell subtype. Top: model proportions of each subtype; Bottom: relative expression of marker genes associated with each cell subtype. G) t-SNE plot of 9,968 B cells, color-coded by cell subtypes. H) Pie plot showing the proportion of different B cell subtypes. I) Complex heatmap of selected marker genes in each B cell subtype. Top: model proportions of each subtype; Bottom: relative expression of marker genes associated with each cell subtype. J) t-SNE plot of 2,093 pDCs, color-coded by cell subtypes. K) Pie plot showing the proportion of different pDC subtypes. L) Violin plot of selected marker genes in each pDC subtype.

### Supplemental Figure S4

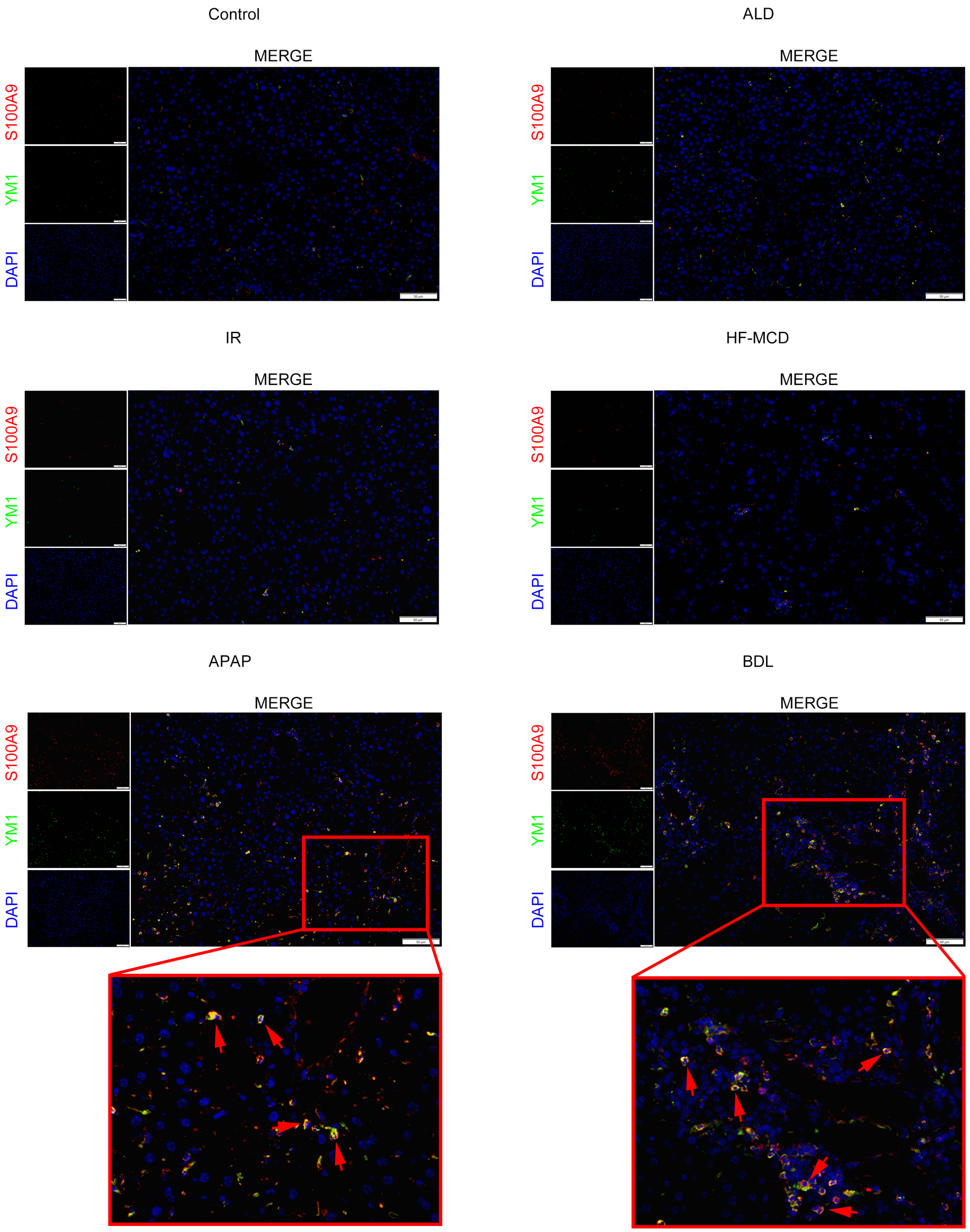

**Supplemental Figure S4.** Immunofluorescence verification experiment for neutrophil subtype. Immunofluorescence staining of Neu3 subtype markers (S100A9 and YM1) in murine livers of all groups. Nuclei were stained using DAPI (blue). Scale bars, 50 μm. Related to figure 3.

### Supplemental Figure S5

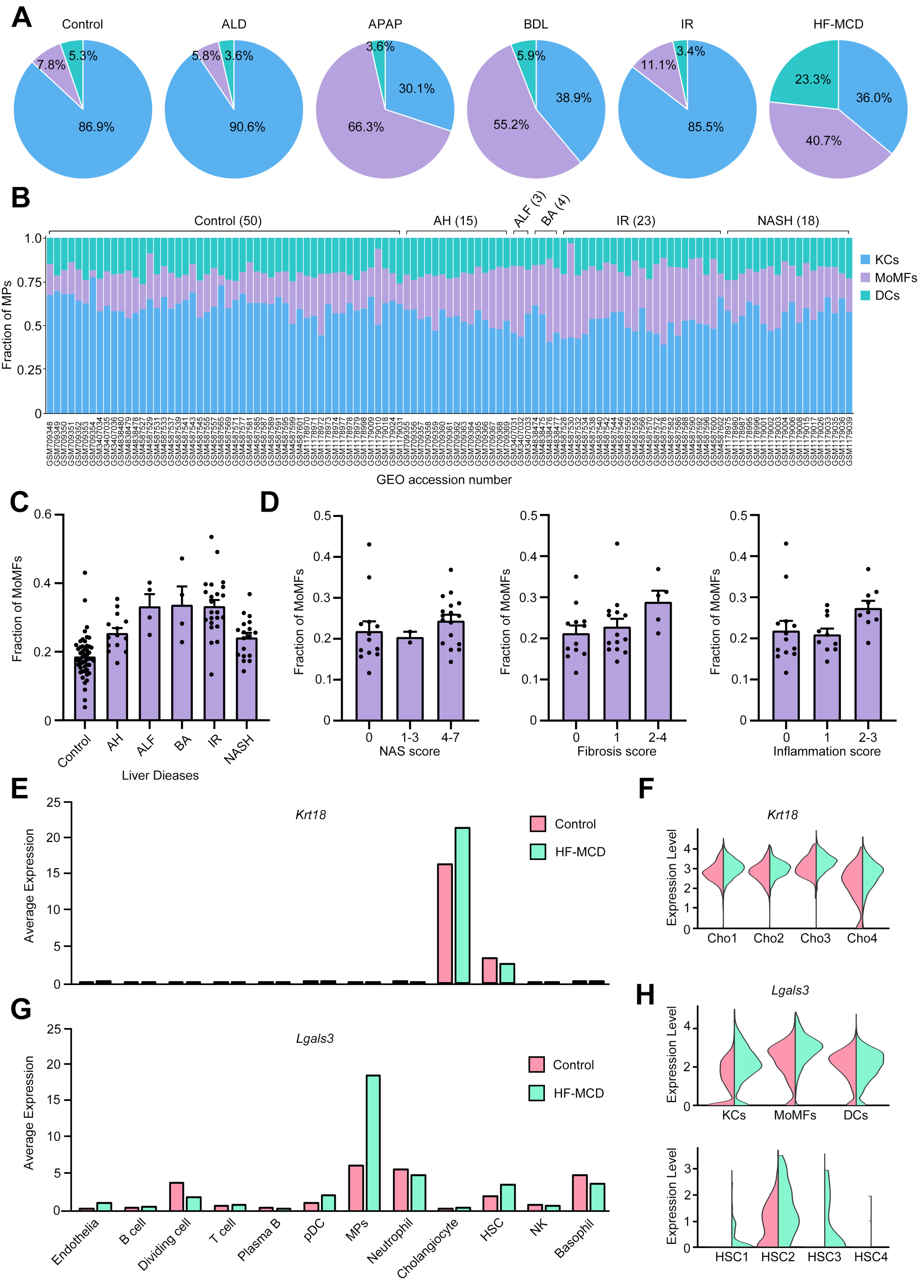

**Supplemental Figure S5.** Combined scRNA-seq data with clinical liver disease samples. A) The MPs composition of different groups in scRNA-seq data. B,C,D) Deconvolution analysis of publicly available microarray/bulk-seq data from different liver disease samples (n = 113). (B) The MPs composition. (C) The fraction of MoMFs in different liver disease samples. (D) Left: frequency of MoMFs in patients with NAS scores of 0, 1-3 and 4-7; Middle: frequency of MoMFs in patients with fibrosis scores of 0, 1 and 2-4; Right: frequency of MoMFs in patients with inflammation scores of 0, 1 and 2-3. E,F) The expression of *Krt18* in different cell types (E) and different cholangiocyte subtypes (F) in control and HF-MCD groups. G,H) The expression of *Lgals3* in different cell types (G) and different MPs (H, top) and HSC (H, bottom) subtypes in control and HF-MCD groups. Related to figure 4.

### Supplemental Figure S6

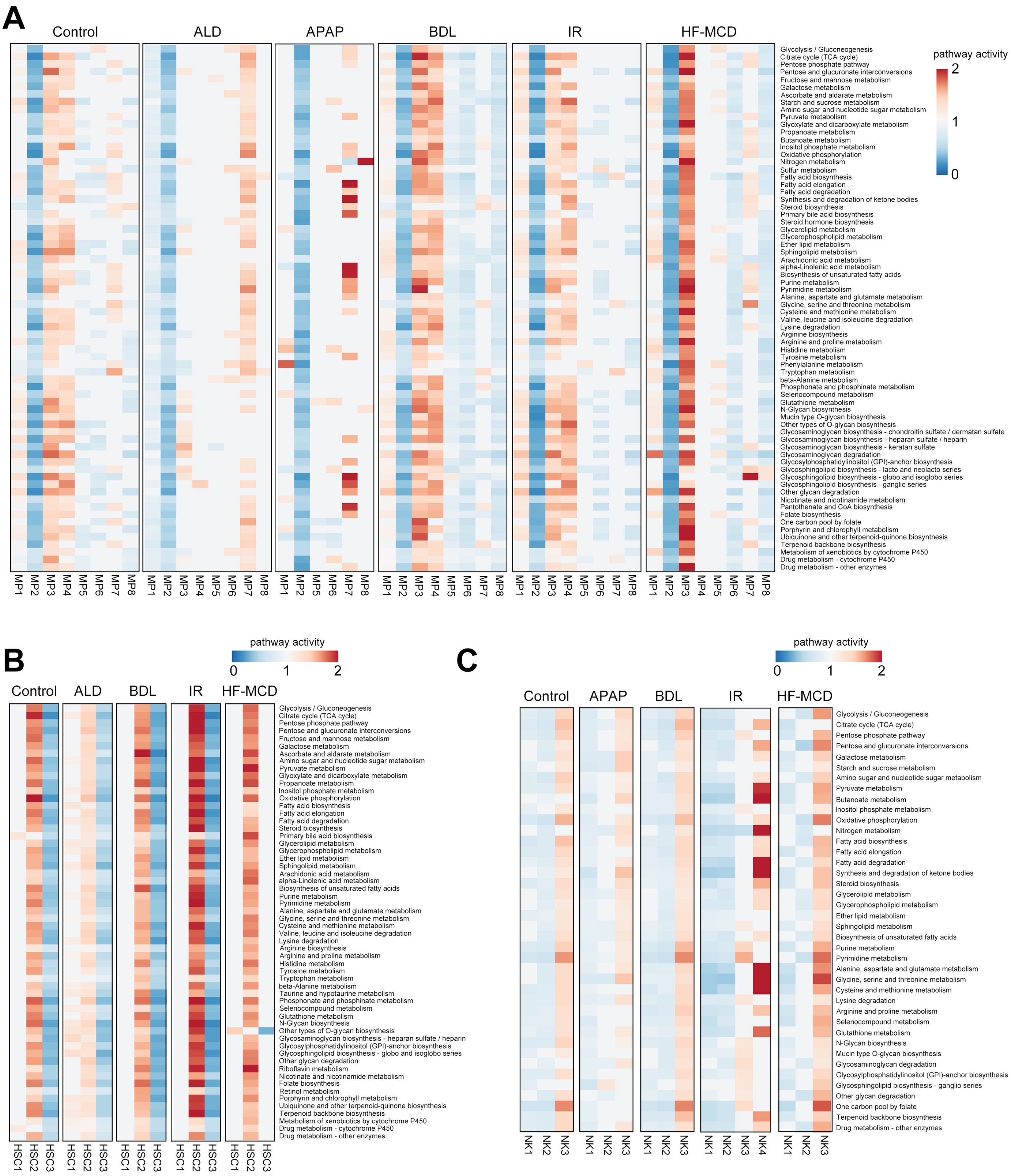

**Supplemental Figure S6.** Disease-specific metabolic reprogramming of each cell subtype in MPs, HSC and NK. A,B,C) Metabolic pathway activities of each MPs (A), HSC (B), and NK (C) subtype in different groups. For MP3, HSC4 and NK4 subtypes, no metabolic pathways were enriched in some groups due to the few cell number. Related to figure 5.

### Supplemental Figure S7

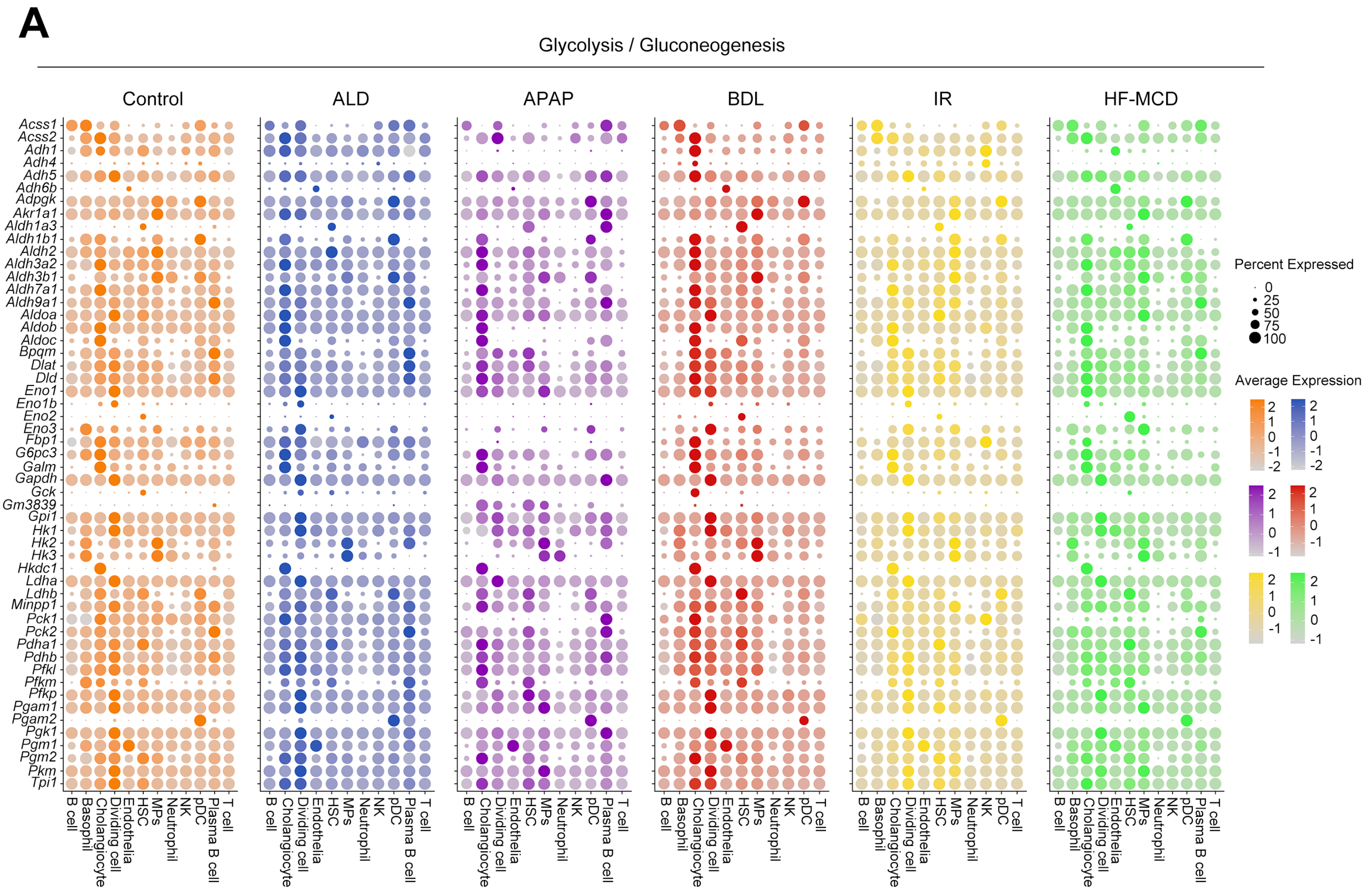


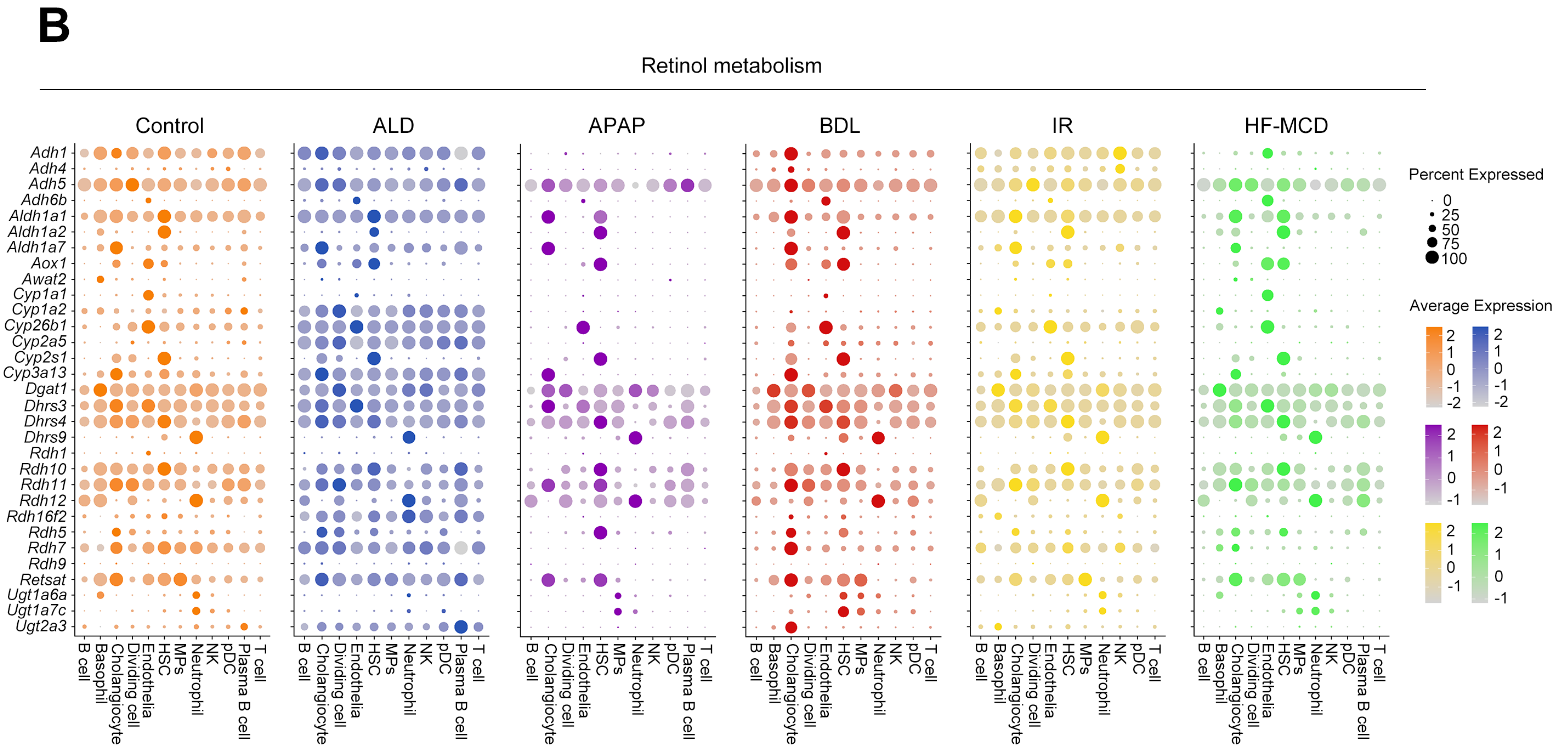

**Supplemental Figure S7.** The expression of specific metabolic pathways related genes of each cell type in different groups. A,B) Dot plot displaying the expression of glycolysis/gluconeogenesis pathway related genes (A) and retinol pathway related genes (B) of each cell type in different models. Size and color of the dot represents express percentage and average expression of gene separately. Related to figure 5.

### Supplemental Figure S8

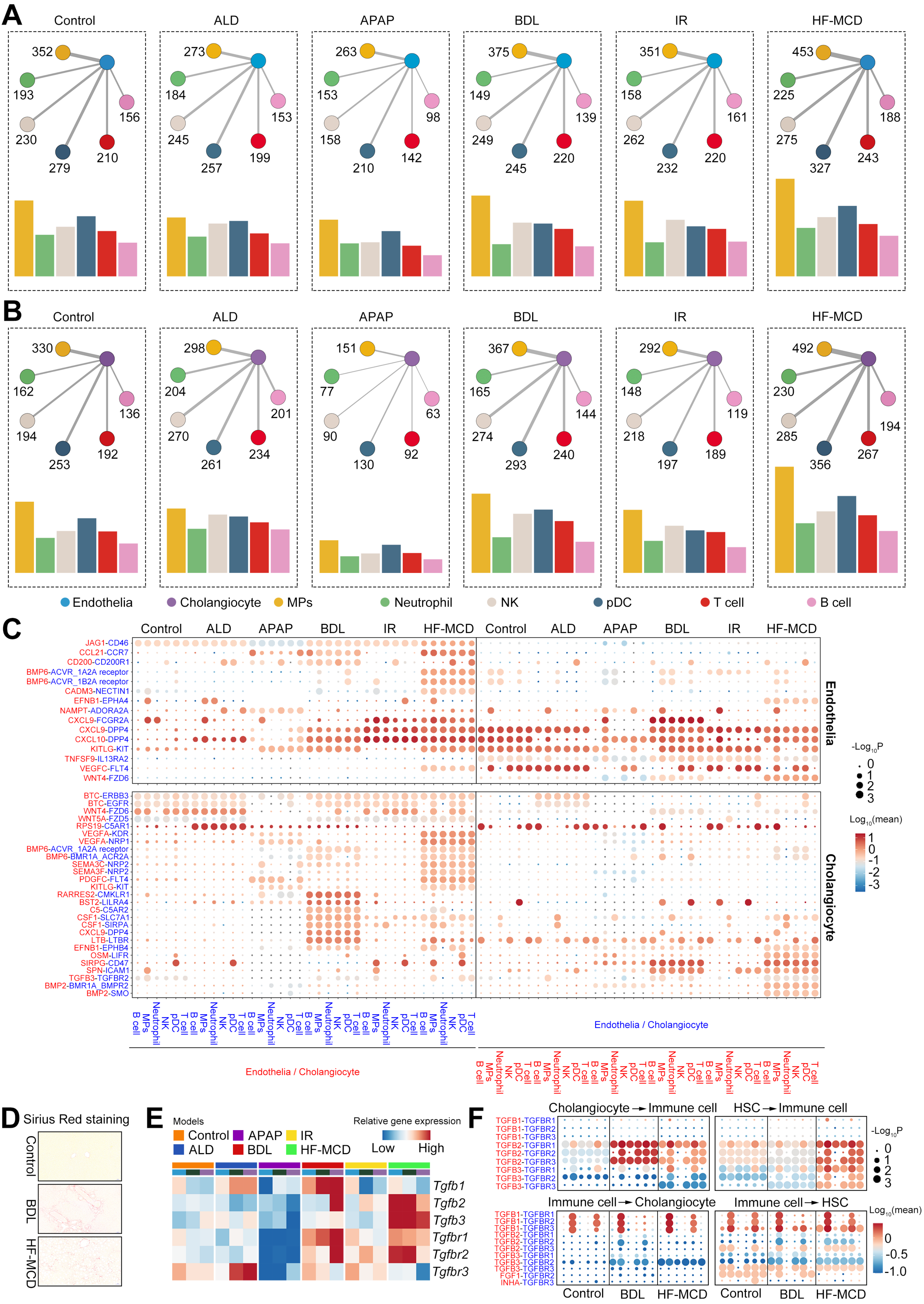

**Supplemental Figure S8.** The CCI network between non-immune cells and immune cells. A,B) The interactions between endothelial cell (A) or cholangiocyte (B) and immune cells. The line thickness is proportional to the number of interactions between two cell types. C) Dot plot displaying the specific ligand-receptor interactions between endothelial cell (top) or cholangiocyte (bottom) and immune cells in different groups. Size of the dot represents statistical significance of the indicated interactions and color of the dot represents the total mean of the individual partner average expression values in the corresponding interacting pairs of cell types. D) Sirius Red staining morphology of murine livers in control, BDL and HF-MCD groups. Scale bars, 50 μm. E) Heatmap showing the relative expression of *Tgfb*- family genes in non-immune cells in different groups F) Dot plot displaying the interactions containing fibrosis-related ligand or receptor genes between cholangiocyte (left) or HSC (right) and immune cells in control, BDL and HF-MCD group. Size of the dot represents statistical significance of the indicated interactions and color of the dot represents the total mean of the individual partner average expression values in the corresponding interacting pairs of cell types. Related to figure 6.

**Supplementary Table S1.** Plasma biochemical parameters of disease mouse models. Related to figure 1A and figure S1A.

| Parameter | ALD model | | APAP model | | BDL model | | IR model | | NASH model | |
|---|---|---|---|---|---|---|---|---|---|---|
| | CON | ALD | CON | APAP | Sham | BDL | CON | IR | NCD | HF-MCD |
| ALT (U/L) | 31.0±5.4 | 200.3±20.5 *** | 37.7±5.6 | 2721.3±1068.3 *** | 17.6±1.9 | 434.9±80.9 *** | 41.0±2.0 | 79.8±14.8 * | 24.8±3.2 | 393.0±61.4 *** |
| AST (U/L) | 100.8±13.5 | 342.7±31.0 *** | 61.8±6.0 | 1581.3±467.7 *** | 41.3±5.6 | 779.5±174.4 ** | 142.5±8.2 | 261.9±39.5 * | 46.0±9.0 | 267.3±39.4 *** |
| TG | 0.45±0.02 | 0.62±0.04 * | / | / | / | / | / | / | / | / |
| CHOL | 1.44±0.09 | 2.22±0.06 *** | / | / | / | / | / | / | / | / |
| TBIL | / | / | / | / | 0.77±0.13 | 299.0±30.3 *** | / | / | / | / |
| ALP | / | / | / | / | 1.57±0.90 | 63.4±10.7 *** | / | / | / | / |

All of the results are expressed as mean ± SEM (n = 3–12/group). CON, control; NCD, normal chow diet; ALD, alcoholic liver disease; APAP, acetaminophen; BDL, bile duct ligation; IR, ischemia-reperfusion; NASH, nonalcoholic steatohepatitis; HF-MCD, high fat-methionine/choline deficient; AST, aspartate aminotransferase; ALT, alanine aminotransferase. TG, triglyceride; CHOL, cholesterol; TBIL, total bilirubin; ALP, alkaline phosphatase. *p < 0.05, **p < 0.01, and ***p < 0.001 vs. CON, NCD, and Sham.
